# Cryptic intronic transcriptional initiation generates efficient endogenous mRNA templates for C9orf72-associated RAN translation

**DOI:** 10.1101/2025.04.27.650888

**Authors:** Shannon L Miller, Katelyn M Green, Bradley Crone, Jessica A Switzenberg, Elizabeth M H Tank, Amy Krans, Karen Jansen-West, Clare M Wieland, Eric W Ji, Leonard Petrucelli, Sami J Barmada, Alan P Boyle, Peter K Todd

## Abstract

Intronic GGGGCC hexanucleotide repeat expansions in *C9orf72* are the most common genetic cause of amyotrophic lateral sclerosis (ALS) and frontotemporal dementia (FTD). Despite its intronic location, this repeat avidly supports synthesis of pathogenic dipeptide repeat (DPR) proteins via repeat-associated non-AUG (RAN) translation. However, the template RNA species that undergoes RAN translation endogenously remains unclear. Using long-read based 5’ RNA ligase-mediated rapid amplification of cDNA ends (5’ Repeat-RLM-RACE), we identified novel C9orf72 transcripts initiating within intron 1 in a C9BAC mouse model, patient-derived iNeurons, and iNeuron-derived polysomes. These cryptic m^7^G-capped mRNAs are at least partially polyadenylated and are more abundant than transcripts derived from intron retention or circular intron lariats. In RAN translation reporter assays, novel intronic template transcripts – even those with short (32 nucleotide) leaders – exhibited robust expression compared to exon-intron and repeat-containing lariat reporters. To assess endogenous lariat repeat RNA contributions to RAN translation, we enhanced endogenous lariat stability by knocking down the lariat debranching enzyme Dbr1. However, this modulation did not impact DPR production in patient-derived iNeurons. These findings identify cryptic, linear, m^7^G-capped intronic-initiating C9orf72 mRNAs as an endogenous template for RAN translation and DPR production, with implications for disease pathogenesis and therapeutic development.

**SIGNIFICANCE STATEMENT:** An intronic GGGGCC repeat expansion in *C9orf72* supports an unusual translational initiation process known as repeat-associated non-AUG (RAN) translation to produce toxic dipeptide repeat (DPR) proteins that contribute to neurodegeneration in ALS and FTD. How an intronic repeat RNA engages with ribosomes to support such translation is unclear. Here we identify a series of novel mRNA transcripts that initiate within the repeat-containing intron to create linear m^7^G-capped templates for RAN translation from GGGGCC repeats. These cryptic mRNAs are present in patient iNeurons, engage with ribosomes, and robustly support RAN translation. This finding has important implications for both our understanding of the mechanism by which RAN translation occurs and on therapeutic development in this currently untreatable class of neurodegenerative disorders.

## INTRODUCTION

The most common genetic cause of both amyotrophic lateral sclerosis (ALS) and frontotemporal dementia (FTD) is a GGGGCC hexanucleotide repeat expansion in the first intron of *C9orf72*, accounting for 30-50% of familial ALS and 7% of sporadic ALS (1–3), as well as up to 25% of familial FTD and 6-8% of sporadic FTD (4). Most healthy individuals have only 2 GGGGCC repeats in C9orf72, but patients diagnosed with ALS/FTD due to C9orf72 mutations (C9ALS/FTD) can harbor hundreds to thousands of repeats (1, 2).

One hallmark of C9ALS/FTD is the presence of dipeptide-repeat-containing proteins (DPRs), which are generated through a non-canonical protein translation initiation mechanism termed repeat-associated non-AUG (RAN) translation. Both sense– and antisense-generated DPRs accumulate within patient brains (5–8). From the sense GGGGCC strand, a poly-glycine-alanine (GA), poly-glycine-proline (GP), and poly-glycine-arginine (GR) containing protein are translated. The antisense CCCCGG strand produces poly-proline-arginine (PR), poly-proline-alanine (PA), and a second poly-GP containing protein, although it is worth noting that the PR and PA reading frames contain an AUG codon that could be used for initiation (9). Overexpression of DPRs across multiple model systems causes neurotoxicity in the absence of repeat RNA or its native sequence context (8, 10–17), suggesting that DPRs may be sufficient for neurodegeneration upon their accumulation.

Multiple groups have utilized reporters to determine how GGGGCC repeats support RAN translation. While there are differences in how the reporters were designed, data from multiple groups suggests the following: RAN translation of the GA reading frame is the most robust (18–24), a CUG codon located 24 nucleotides upstream from the repeat is important for efficient initiation of translation in the GA reading frame (18, 19, 21, 25–29), GGGGCC repeats are most robustly translated from an m^7^G capped mRNA (18, 19, 21, 22, 30), and *C9orf72*-associated RAN translation (C9RAN) is upregulated when the integrated stress response is activated (18, 20–22, 31).

While these reporter assays have yielded valuable insights into disease, the exact RNA template that undergoes RAN translation in patients remains unclear. Typically, introns are spliced from mature mRNA transcripts as a circularized intron lariats and then rapidly turned over via linearization by the lariat debranching enzyme Dbr1. Once linearized, they are subsequent degraded by nuclear exonucleases, precluding their export to the cytoplasm and ability to initiate translation through interaction with ribosomes.

Understanding the template for C9RAN translation and how the intronic GGGGCC repeat in *C9orf72* escapes this fate are areas of active study **(Figure 1A)**. The GGGGCC repeat could theoretically escape degradation and exit to the cytoplasm by impaired splicing, by aberrantly stabilizing a correctly spliced intron, or through inclusion within transcripts with novel 5’ or 3’ ends (28). A splicing failure would lead to retention of the repeat and first intron within mature C9orf72 mRNA – allowing for its export to the cytoplasm and engagement of translation machinery (32, 33). Alternatively, if the spliceosome successfully removed the intron containing the GGGGCC repeat, then the robust secondary structure formed by the repetitive GC rich sequence could aberrantly stabilize the lariat. Spliced introns have been previously reported in the cytoplasm of cells (34), and in C9ALS/FTD, C9 intron lariat RNA persists in the cytoplasm both in cells transfected with a splice-capable reporter and in patient-derived fibroblasts as measured by smFISH (35). Introns have historically been understood to be non-coding, but some circular RNAs, which arise from back splicing events, are translated in a cap-independent manner (36, 37).

**Figure 1:**
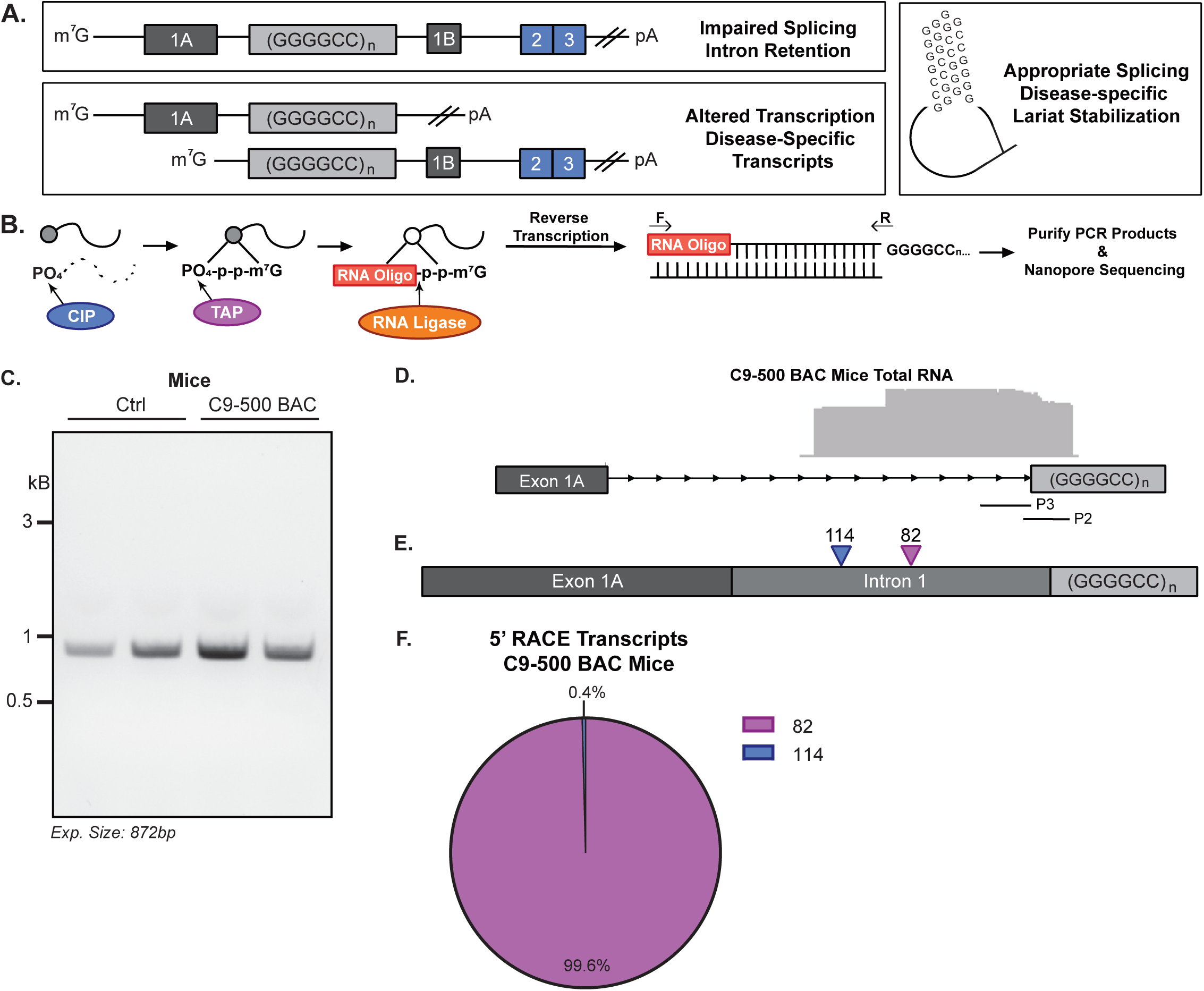
5’ RACE reveals intronic initiation of C9orf72 repeat RNAs in C9BAC mouse model. A) Schematic of potential GGGGCC repeat containing mRNA templates for RAN translation in C9ALS/FTD. B) Schematic of 5’ RACE-RLM experimental procedure. C) Agarose gel of murine Beta-actin 5’ RACE PCR in C9-500 BAC and control mice, *n=2.* D) Aligned IGV plots using primers 2 and 3 in cerebellum of C9-500 BAC mice, *n=3*. E) Annotated schematic of the most commonly identified 5’ RACE products using primers 2 and 3 in cerebellum of C9-500 BAC mice, *n=3*. F) Pie chart of most commonly identified 5’ RACE products using primers 2 and 3 in cerebellum of C9-500 BAC mice, *n=3/group*.

Finally, it is possible that the GGGGCC repeat interferes with proper *C9orf72* transcription, generating transcripts that lack the canonical 5’ or 3’ splice sites to place the repeat in a non-intronic context. This scenario could arise from altered transcription initiation from previously unmapped 5’ start sites or by premature transcription termination, leading to 3’ truncated mRNA (38).

Defining the GGGGCC repeat-containing mRNA transcript template that supports RAN translation is critical both for our understanding of this enigmatic process and for development of therapies for C9ALS/FTD. Here, we describe a set of novel 5’ m^7^G-capped C9orf72 RNAs that are derived from cryptic initiation within intron 1 of the C9 locus. These cryptic transcripts are selectively present in C9BAC-500 mice compared to controls and are also observed in patient-derived fibroblasts, patient-derived iNeurons, and iNeuron-derived polysomes. Enrichment of poly-adenylated RNA captured these novel transcripts as well as additional transcripts with initiation sites in or above exon 1, indicating that a portion of these novel transcripts are poly-adenylated and some likely derive from intron retention, respectively. Using reporters, we find that these novel 5’ intronic-initiating transcripts are translated much more robustly than exon-intron or lariat reporters with the same sized repeats. Moreover, efforts to enhance translation from endogenous C9 intron lariat RNA through lariat stabilization did not impact DPR production in patient-derived iNeurons. Taken together, these data define a novel native template for RAN translation from the GGGGCC repeat in *C9orf72*, with implications for the mechanism by which this translation occurs and for therapies aimed at mitigating this process.

## RESULTS

### 5’ RACE reveals novel intronic-initiated C9orf72 repeat-containing mRNAs in C9BAC mouse model

We used a BAC transgenic mouse model engineered to express full-length human *C9orf72* with 500 GGGGCC repeats in intron 1 as well as human surrounding sequence (C9-500 BAC) (39). This mouse recapitulates key molecular and pathological features of C9ALS/FTD, including production of sense and antisense repeat containing RNA species and generation of polyGA, polyGP, and polyGR DPRs. As the cerebellum of these mice are particularly rich polyGA and polyGP aggregates (39, 40), we isolated total RNA from cerebellar homogenates from 6-week-old mice and performed 5’ RNA ligase mediated-rapid amplification of cDNA ends (5’ Repeat-RLM-RACE) to identify the 5’ end of transcribed *C9orf72* GGGGCC repeat-containing mRNAs (**Figure 1B**). cDNA was synthesized using random hexamers and spiked in (CCCCGG)_4_ to increase the probability of the reverse transcriptase making it through the repeat, which is known to impair PCR (2). Unlike traditional 5’ RACE, we developed a long-read nanopore sequencing platform of our PCR amplicons rather than TOPO cloning to create a high-throughput analysis pipeline, allowing us to better capture both high and low-frequency transcripts and to more accurately determine the frequency of different products. Control RACE PCR reactions using primers specific for murine beta-actin and the 5’ ligated RNA oligo (**Table 1**) revealed a single band of the expected size on an agarose gel, confirming that all steps of the reaction were successful (**Figure 1C**).

**Table 1:**
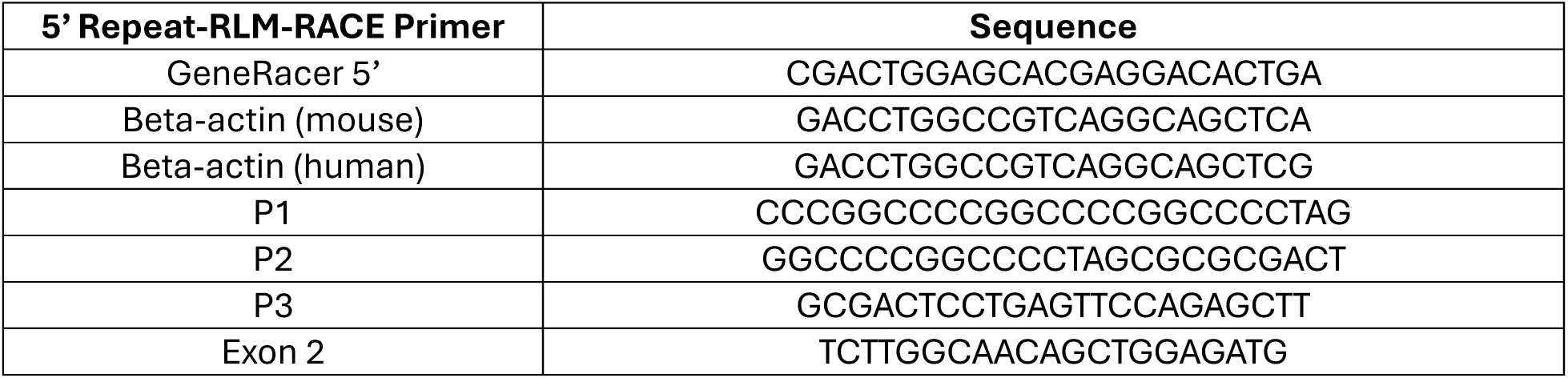
5’ Repeat-RLM-RACE Reverse Primer Sequences.

**Table 2:**
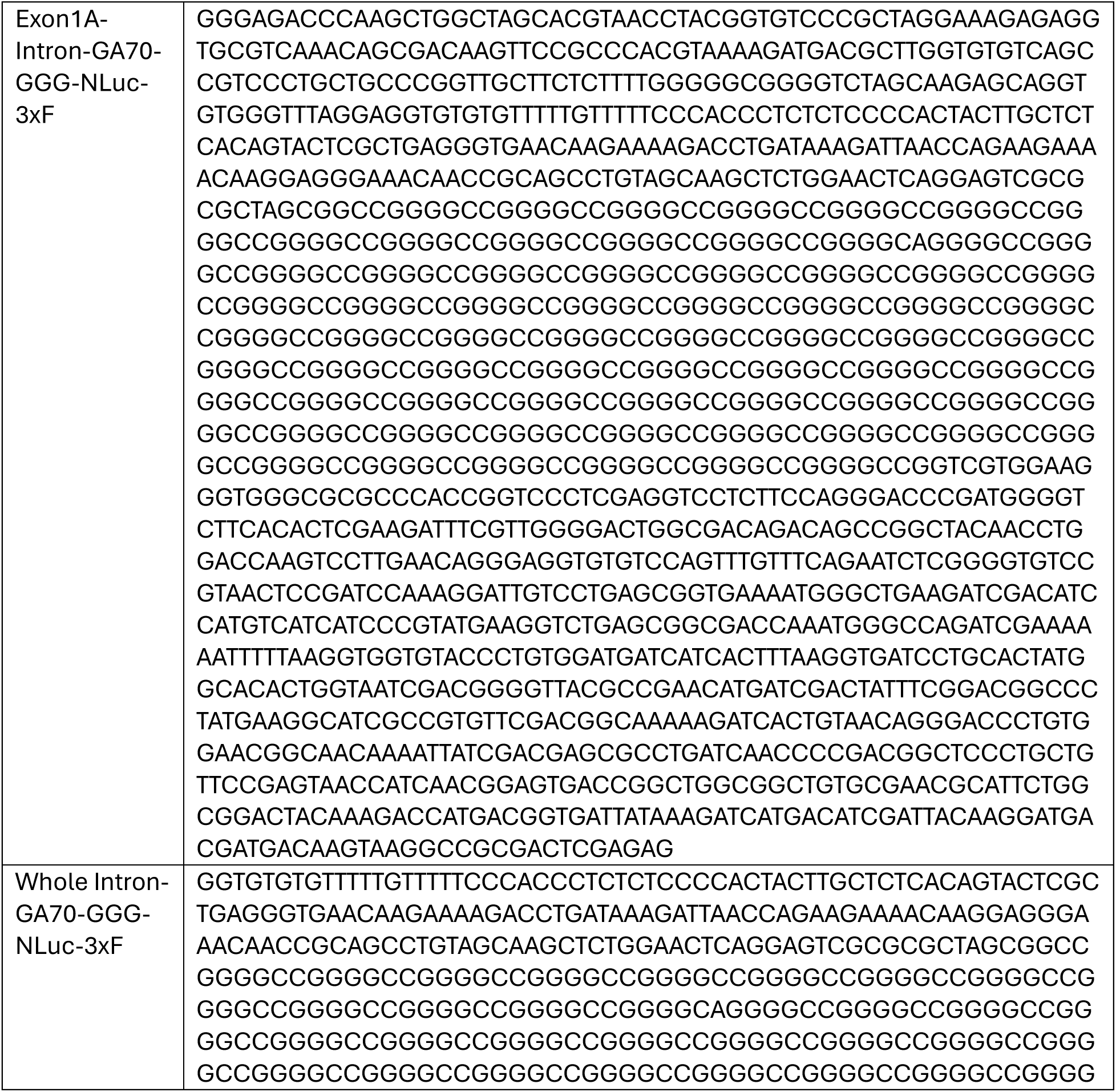
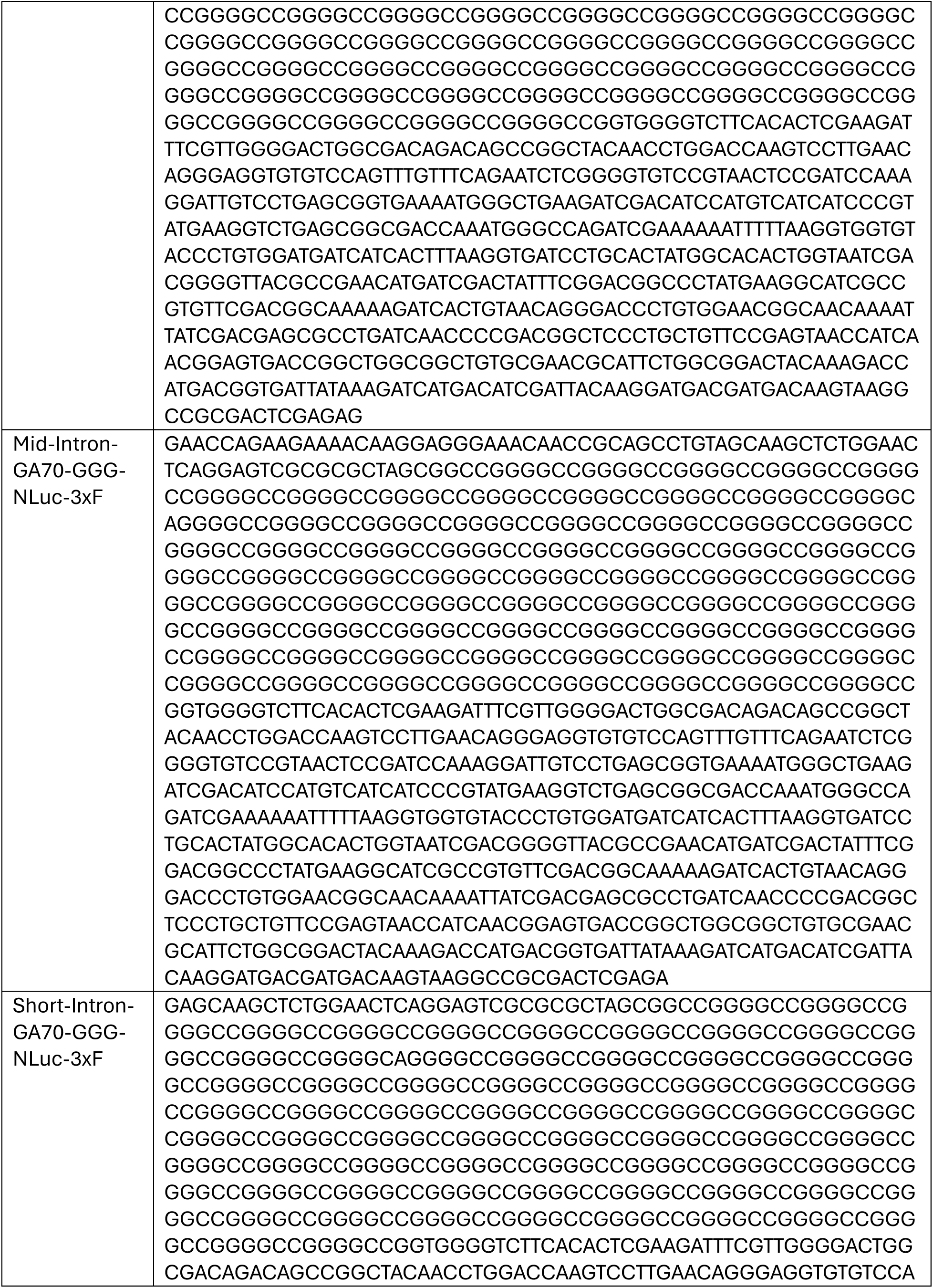
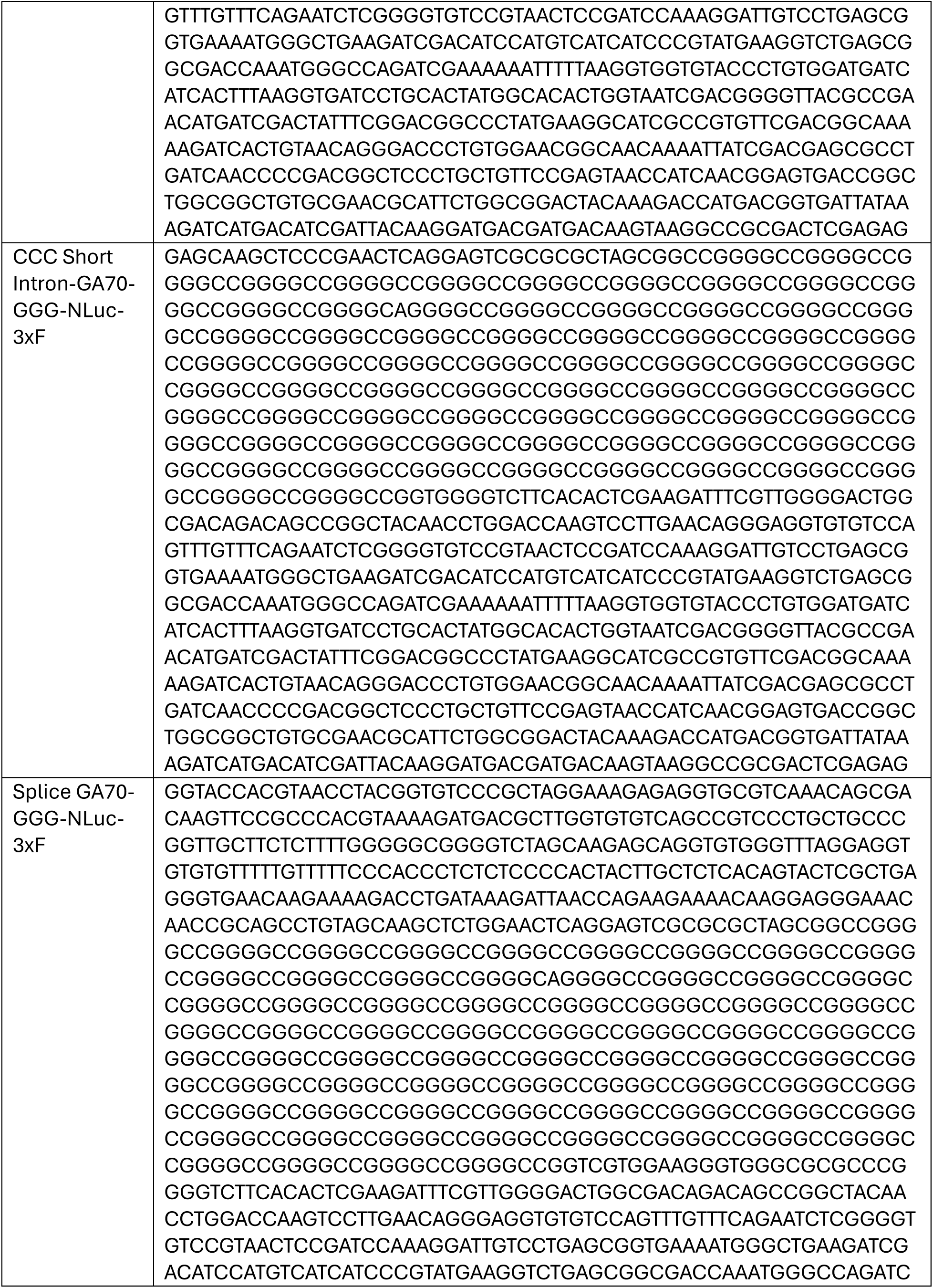
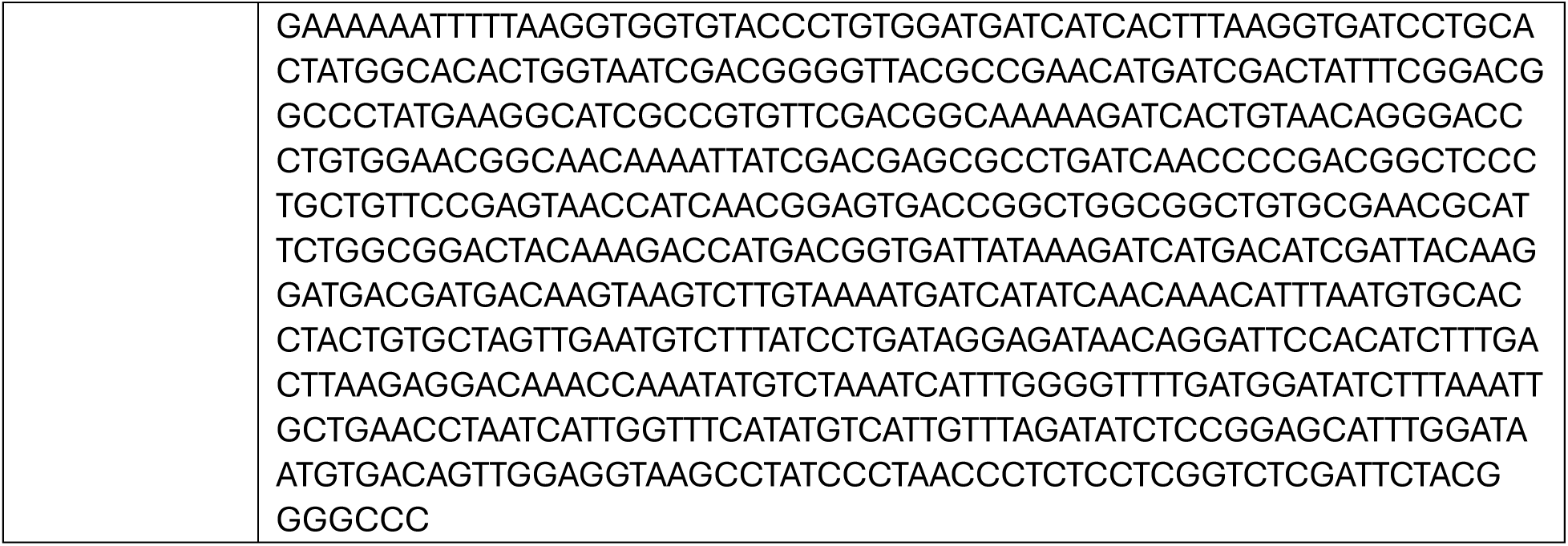
Reporter Sequences.

**Table 3:**
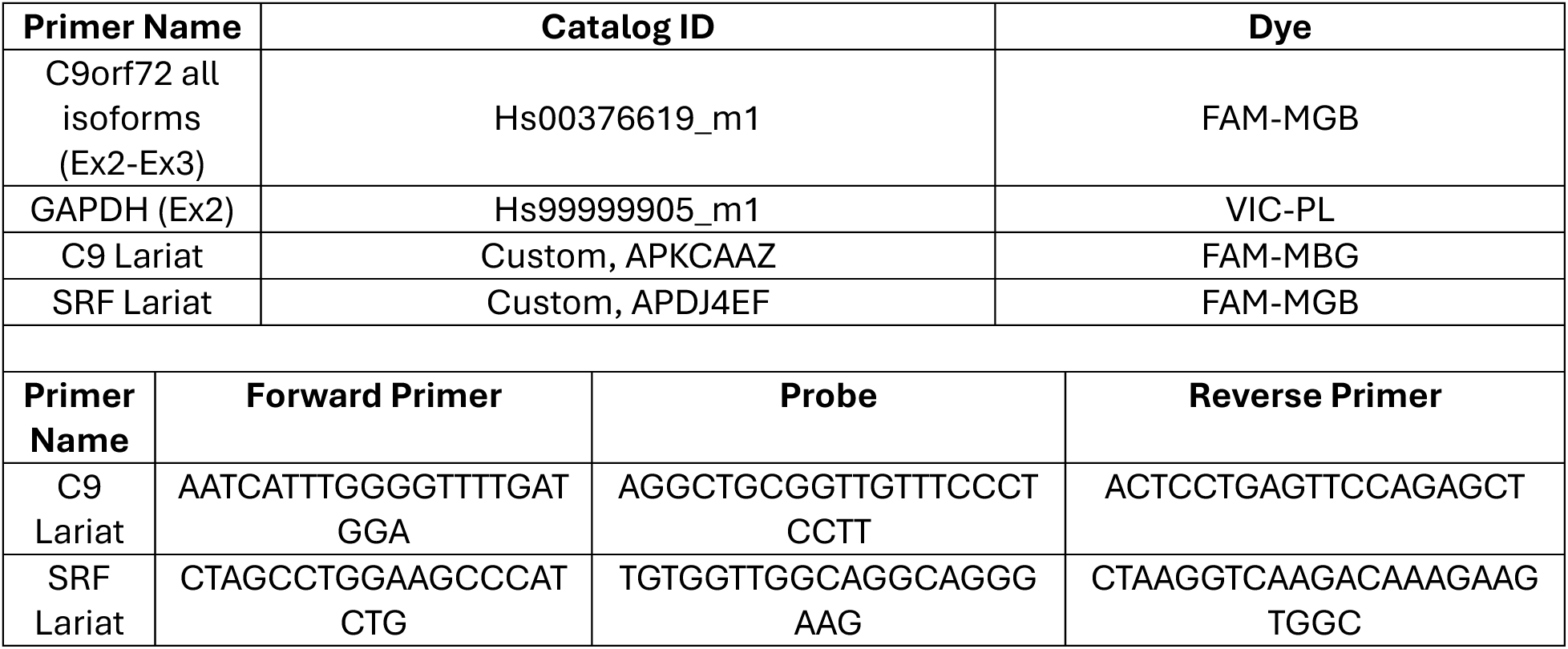
qRT-PCR Primers.

**Table 4:** siRNA Sequence.

| siRNA | Source | Catalog Number/siRNA ID |
| --- | --- | --- |
| Dbr1 | Thermo silencer select | s27586 |
| NT control | Thermo silencer select | 4390843 |

**Table 5:**
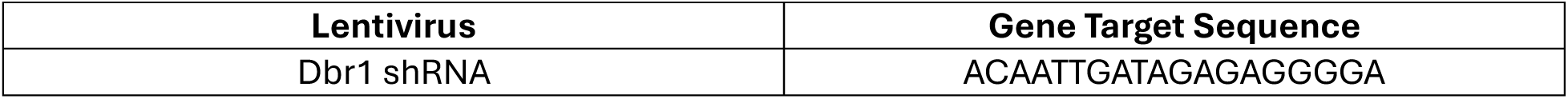
Lentiviral shRNA Target Sequence.

We initially used three RACE primers specific to intron 1 of *C9orf72* (P1, P2, and P3, **Table 1**), but found that the lack of sequence specificity of P1 did not allow for C9-specific products to be identified. However, using P2 and P3, we identified a novel 5’ end transcript that initiated within intron 1, 82 nucleotides (nt) upstream of the repeat in C9-500 BAC mice (**Figure 1D-F**). At very low frequency (<1%), one additional product was identified, initiating at 114nt of the repeat (**Figure 1D-F**). Both of the identified transcripts contained the CUG codon located 24nt upstream of the repeat thought to drive translation of the repeats in the polyGA frame (18, 19, 21, 25). These products were not reliably detected from control mice, as P1, P2, and P3 do not have significant sequence homology with mouse *C9orf72* to allow for binding (**Supplemental Figure 1A**). Of note, we did not identify products mapping to known transcription start sites in exon 1A in our mouse samples despite PCR cycling parameters that would allow sufficient time for these products to be amplified, suggesting that most repeat-containing mRNAs do not arise from intron retention.

### 5’ RACE in C9 patient-derived cells identifies additional intronic-initiating C9orf72 RNAs

In patient-derived cells, we initially utilized a traditional low-throughput 5’ RACE protocol to determine whether novel 5’ end products are also present in human cells. RNA was harvested from one control and two C9ALS/FTD fibroblast lines, and PCR products from P1, P2, and P3 amplification reactions were run on an agarose gel. Bands were excised, gel purified, TOPO cloned, and Sanger sequenced. Using P2 and P3, we identified a short product in both C9ALS/FTD and control fibroblasts initiating 32nt upstream of the repeat and only 8nt 5’ to the CUG near-cognate initiation codon (**Supplemental Figure 1B**). This mRNA initiates at a –1T, +1A motif that is enriched at transcription start sites. However, only fourteen total clones mapped to *C9orf72* over multiple reactions. While more clones came from C9ALS/FTD fibroblast lines, the presence of this product in at least one control line suggests that it is not disease specific. We identified no *C9orf72*-specific products with the P1 primer from C9ALS/FTD or control fibroblasts.

To determine whether these novel 5’ end transcripts undergo RAN translation in disease-relevant cells, we generated iNeurons from human induced pluripotent stem cells (iPSCs) containing an NGN2-cassette from one control and one C9ALS/FTD line. Utilizing iNeuron lysates, we performed polysome profiling and captured RNA from 80S and polysome fractions to use as our input for 5’ RACE (**Figure 2A**). Sequencing of clones that utilized the P1 primer did not yield any C9orf72-specific hits, but clones derived from the P3 primer identified sixteen C9orf72 hits that mapped to the 32nt product seen in fibroblasts (**Figure 2B**). We also observed a rare, longer C9orf72 isoform in our C9ALS/FTD iNeuron line initiating near the mid-point of intron 1 (**Figure 2B**). Thus, novel, short, 5’-capped C9orf72 transcripts are present within fibroblasts and actively translating fractions of iNeuron RNA, suggesting these species could contribute to DPR production.

**Figure 2:**
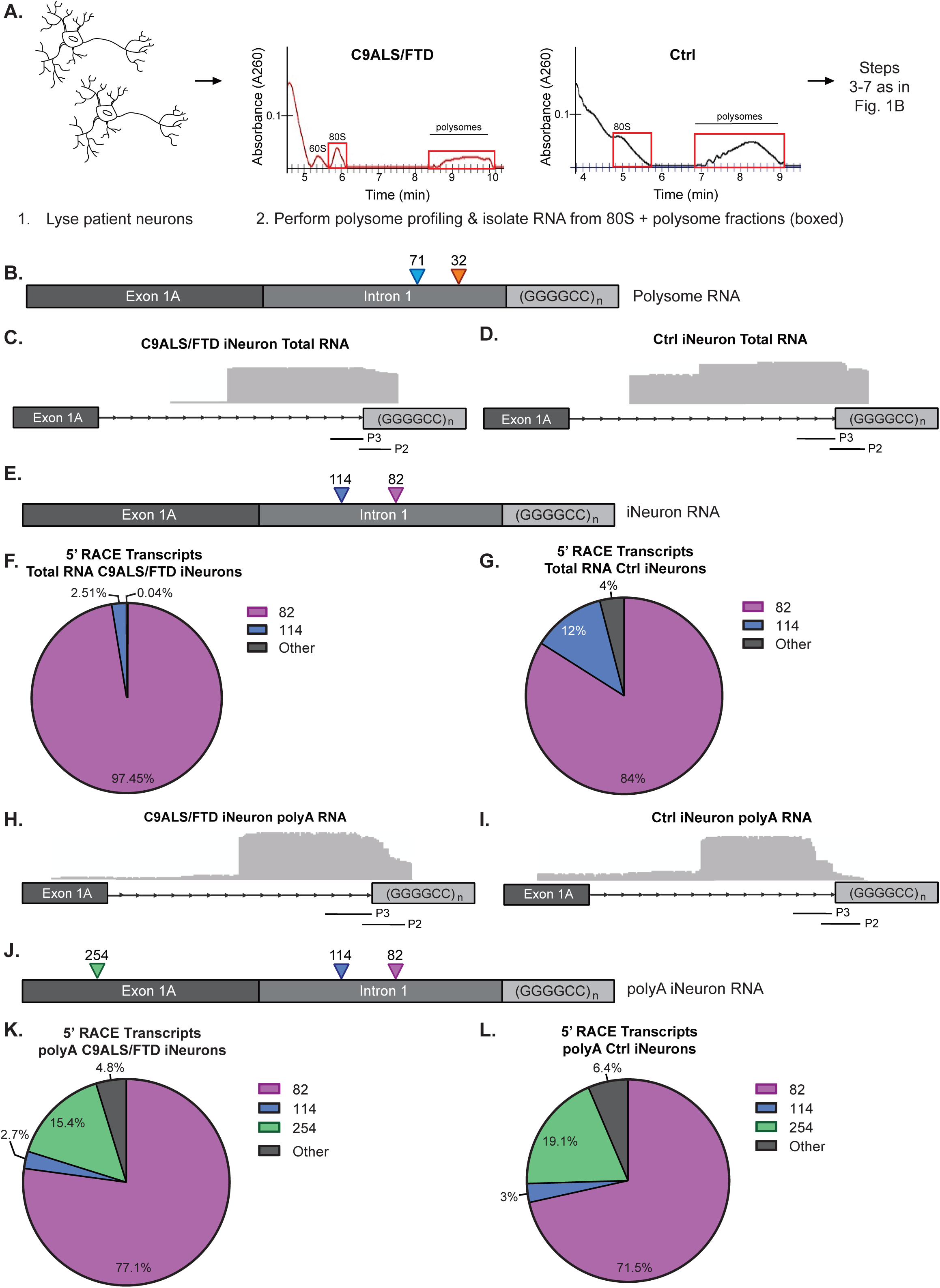
5’ RACE reveals intronic-initiating C9orf72 repeat RNAs in C9ALS/FTD patient-derived iNeurons. A) Schematic of 5′ RLM-RACE experimental flow for RNA isolated from control and C9orf72 patient-derived iNeuron 80S and polysome fractions. Polysome profiles shown were obtained from ∼90 μg total RNA isolated from C9ALS/FTD iNeurons (left) and control iNeurons (right), with 80S and polysome fractions labeled. B) Annotated schematic of the most commonly identified 5’ RACE products using primers 1, 2 and 3 in polysome-associated RNA from C9ALS/FTD and control iNeurons, *n=1/group.* C) Aligned IGV plots using primers 2 and 3 in total RNA from C9ALS/FTD iNeuron lines, *n=3.* D) Aligned IGV plots using primers 2 and 3 in total RNA from control iNeuron lines, *n=3*. E) Annotated schematic of the most commonly identified 5’ RACE products using primers 2 and 3 in total RNA from C9ALS/FTD and control iNeuron lines, *n=3/group.* F) Frequency of most commonly identified 5’ RACE products using primers 2 and 3 in total RNA from C9ALS/FTD iNeuron lines, *n=3/group*. G) Frequency of most commonly identified 5’ RACE products using primers 2 and 3 in total RNA from control iNeuron lines, *n=3/group.* H) Aligned IGV plots using primers 2 and 3 in polyA RNA from C9ALS/FTD iNeurons, *n=3.* I) Aligned IGV plots using primers 2 and 3 in polyA RNA from control iNeurons, *n=3.* J) Annotated schematic of the most commonly identified 5’ RACE products identified using primers 2 and 3 in polyA-captured RNA from C9ALS/FTD and control iNeuron lines, *n=3/group.* K) Frequency of most commonly identified 5’ RACE products using primers 2 and 3 in polyA-captured RNA from C9ALS/FTD iNeuron lines, *n=3/group.* L) Frequency of most commonly identified 5’ RACE products using primers 2 and 3 in polyA-captured RNA from control iNeuron lines, *n=3/group*.

Due to the small number of clones derived with this low throughput Sanger sequencing based method, we leveraged our higher throughput pipeline with nanopore sequencing on a larger cohort of iNeuron lines as was done with the C9-500 BAC mice. RNA from three control and three C9ALS/FTD iNeuron lines were harvested at day 10 post differentiation and processed for 5’ RACE. The presence of a single band of the expected size using control primers complementary to human beta-actin and 5’ RNA oligo primers on an agarose gel confirmed all steps of the reaction were successful (**Supplemental Figure 1C**). In addition, nanopore sequencing of products generated using a reverse primer in exon 2 showed 5’ ends which mapped to exon 1B and 1A, respectively (**Supplemental Fig. 1D**), demonstrating the presence of correctly spliced, mature RNA in both C9ALS/FTD and control iNeurons and suggesting that the RACE process from the C9 locus is unimpeded by the repeat. Nanopore sequencing of PCR products from P2 and P3 identified the same 82nt product seen in C9-500 BAC mice which represented ∼97% and 85% of the total products in C9ALS/FTD and control iNeurons, respectively, as well as the 114nt product, representing 2.5% and 12% of the total products, respectively (**Figure 2C-G**) As in the C9-500 BAC mice, we did not identify repeat-RLM-RACE reads from total RNA in C9ALS/FTD iNeurons mapping to exon 1A, suggesting that most repeat-containing mRNAs in human iNeurons do not arise from intron retention. However, a small percentage (4%) of reads from control iNeurons mapped 5’ to exon 1A (**Figure 2G**).

To assess whether these novel intronic initiating C9orf72 RNAs were polyadenylated, we performed polyA-capture using oligo-dT beads prior to performing the nanopore-based repeat-RLM-RACE protocol on the same control and C9ALS/FTD iNeuron lines we harvested total RNA from. Two products in the polyA-captured RNA matched those seen in C9-500 BAC mice and total RNA from C9ALS/FTD and control iNeurons (82nt and 114nt) (**Figure 2H-L**). The recurrence of these products in our polyA-captured 5’ RACE dataset suggests that at least some of these novel intronic initiating products are polyadenylated. Interestingly, ∼15-20% of polyA-captured products mapped back to exon 1A, suggesting at least some RNAs retain intron 1 and that these RNAs are largely polyadenylated (**Figure 2H-L**). Thus, across multiple cell types and systems, we identified novel, intronic-initiating C9orf72 RNAs, and demonstrate that at least a percentage of these RNAs are polyadenylated.

### RAN translation of intron-initiated mRNAs exhibit cap dependence and CUG codon usage

To understand more about the novel 5’ end C9orf72 RNAs we identified by 5’ RACE, we generated a series of nanoluciferase (NLuc) reporters to assess their translation efficiency across multiple systems (**Figure 3A**). Briefly, the AUG start codon of NLuc was mutated to GGG, precluding translation, as previously shown (18, 41). Upstream of GGG-NLuc we inserted 70 GGGGCC repeats in the polyGA (+0), polyGP (+1), or polyGR (+2) reading frame. The 5’ ends we identified through our RACE studies were then inserted upstream of the repeats and immediately downstream of the T7 promoter, so no additional vector sequence was present in *in vitro* transcribed RNAs from these plasmid reporters. We also generated a “whole intron” reporter encompassing all 162 nucleotides of intron 1 to compare to the novel, shorter within-intron initiating RNAs (“short intron” and “mid intron”, respectively). Of note, all reporters contained the CUG start codon located 24 nucleotides upstream of the GGGGCC repeat.

**Figure 3:**
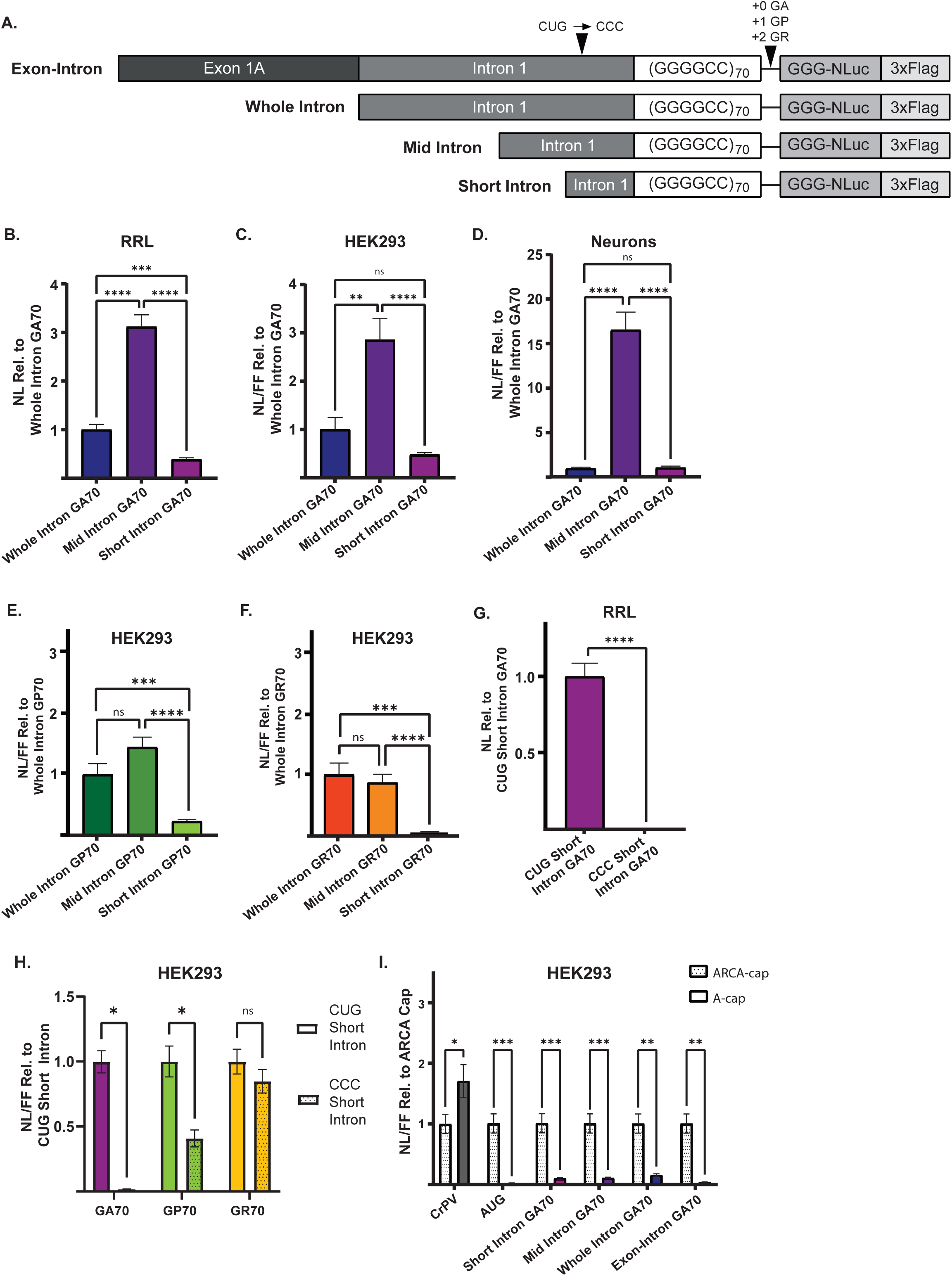
Novel intronic-initiating reporter RNAs are translated in a cap dependent manner and more robustly than intron inclusion RNAs. A) Schematic of linear nanoluciferase (NLuc) reporter mRNAs, including mRNA with short (32nt) leader. B) Expression of short, mid and whole intron GA70 reporters in rabbit reticulocyte lysate (RRL), expressed as NLuc (NL) relative to whole intron GA70, *n=9.* C) Expression of short, mid and whole intron GA70 reporters in HEK293 cells, expressed as NL/FF relative to whole intron GA70, *n=9.* D) Expression of short, mid and whole intron GA70 reporters in rat hippocampal neurons, expressed as NL/FF relative to whole intron GA70, *n=15.* E) Expression of short, mid and whole intron GP70 reporters in HEK293 cells, expressed as NL/FF relative to whole intron GP70, *n=9.* F) Expression of short, mid and whole intron GR70 reporters in HEK293 cells, expressed as NL/FF relative to whole intron GR70, *n=9.* G) Expression of short intron GA70 reporters in RRL, expressed as NL relative to CUG short intron GA70. polyGA frame CUG was mutated to CCC, *n=9.* H) Expression of short intron GA70, GP70, and GR70 reporters in HEK293 cells, expressed relative to CUG short intron for each frame. polyGA frame CUG was mutated to CCC, *n=9.* I) Expression of ARCA m^7^G-capped and A-capped short intron, mid intron, whole intron and exon-intron GA70 reporters in HEK293 cells, expressed as the ratio of NL/FF signal in A-capped reporters to m^7^G-capped reporters, *n=9.* Graphs represent mean ± SEM, \**p* < 0.05, *** *p* < 0.001, **** *p* < 0.0001. 3xF = 3x FLAG tag; GA = glycine–alanine; GP = glycine–proline; GR = glycine– arginine; CrPV = cricket paralysis virus. (B-F) Brown-Forsythe one-way ANOVA with Welch’s unpaired t-test multiple comparison correction, (G) Two-tailed student’s T-test with Welch’s correction, (H-J) Multiple Holm-Šídák two-tailed student’s T-test with Welch’s correction.

Using *in vitro* transcribed RNAs from these reporters, we observed that the mid intron reporter, which corresponds to the most common product identified by 5’ RACE, was expressed more robustly in the GA reading frame than the whole intron or the short intron RNA reporter *in vitro* in rabbit reticulocyte lysate (RRL), HEK293 cells, and rat hippocampal neurons (**Figure 3B-D**). The mid intron RNA reporter was also expressed to the highest extent in the GP and GR reading frames both *in vitro* and in rat hippocampal neurons (**Supplemental Fig. 2A-C**). In HEK293 cells, while the mid intron reporter was expressed more robustly than the short intron RNA reporter in the polyGP and polyGR frames, expression was not significantly increased from the whole intron RNA reporter (**Figure 3E-F)**. Next, we assessed whether the CUG codon 24 nucleotides upstream of the repeat was obligate for RAN translation from these shorter leaders. Typically, there is a 60-nucleotide region at the end of 5’ m^7^G capped mRNA transcripts that is not permissive for initiation due to eIF4F binding (42). Despite being located only eight nucleotides downstream from the 5’ cap, the CUG codon was important for polyGA production from the short intron reporter in both RRL and HEK293 cells, as mutation to a CCC completely abolished translation (**Figure 3G-H**). The loss of the CUG codon had a more modest inhibitory effect on translation from the polyGP in both systems (**Figure 3H, Supplemental Figure 2D**), while its loss in the polyGR frame was variable across systems (**Figure 3H, Supplemental Figure 2E)**, consistent with data suggesting frameshifting may occur within the repeat (18, 19).

The 5’ RACE protocol selectively enriches for 5’ m^7^G-capped mRNAs. We therefore assessed whether the translation of these intronic initiated transcripts exhibited 5’ m^7^G-cap dependence. RNAs were *in vitro* transcribed with either a 5’ m^7^G-or an A-cap analog that cannot recruit the cap binding initiating factor eIF4E but protects the mRNA from degradation. As expected, translation from Cricket paralysis virus (CrPV), which utilizes a cap-independent internal ribosome entry site (IRES) translation mechanism (43), was not impaired when A-capped, while an AUG-initiated NLuc reporter was strongly cap-dependent (**Figure 3I, Supplemental Figure 3A-E**). In RRL and HEK293 cells, the lack of a canonical 5’ m^7^G cap dramatically reduced translation of our novel 5’ end reporter RNAs in all reading frames and at all intron leader lengths, including the 32-nucleotide leader short intron mRNA (**Figure 3I, Supplemental Figure 3A-E**). An intron retention reporter RNA also demonstrated strong cap dependence (**Figure 3I, Supplemental Figure 3A**). In conclusion, our findings demonstrate that intron-initiated *C9orf72* RNAs identified through 5’ RACE are efficiently translated, with the mid intron products corresponding to enhanced expression and even the short leader construct demonstrating a strong dependence on the CUG near-cognate start codon and m^7^G cap, highlighting the critical role of these elements in C9RAN translation.

### Intron-initiated mRNAs are an efficient template for RAN translation

Prior studies have suggested that the major template for RAN translation is either a linear mRNA containing a retained intron (32, 33), or a spliced, circular C9 intron lariat that is exported from the nucleus to the cytoplasm (35). To directly test the efficiency of translation from these potential templates, we generated constructs that included both Exon1A and all intronic sequence 5’ to the repeat, followed by 70 GGGGCC repeats in the polyGA frame upstream of NLuc. This construct contains a predicted AUG initiated uORF that would terminate 79 bases from the start of the repeat (19, 44). We next modified this construct to directly assess a role for C9 intron lariat-based RAN translation by expressing a NLuc reporter plasmid with 70 GGGGCC repeats in the polyGA frame positioned between the 5’ and 3’ ends of intron 1, flanked by Exon 1A upstream and roughly 50 nucleotides of Exon 2 fused with a V5 c-terminal tag, excluding the canonical AUG start codon downstream (lariat GA70) (**Figure 4A**). When transfected into cells, this reporter undergoes splicing to create a circular C9 intron lariat containing the repeat (**Figure 4A**). To confirm the presence of a spliced circular C9 intron lariat, we designed primers to amplify the region of the lariat that crosses the branch point. Importantly, this sequence is only present if the intron is circularized, and sequencing of HEK293 cells expressing lariat GA70 confirmed our reporter generated the anticipated product (**Supplemental Figure 4A**).

**Figure 4:**
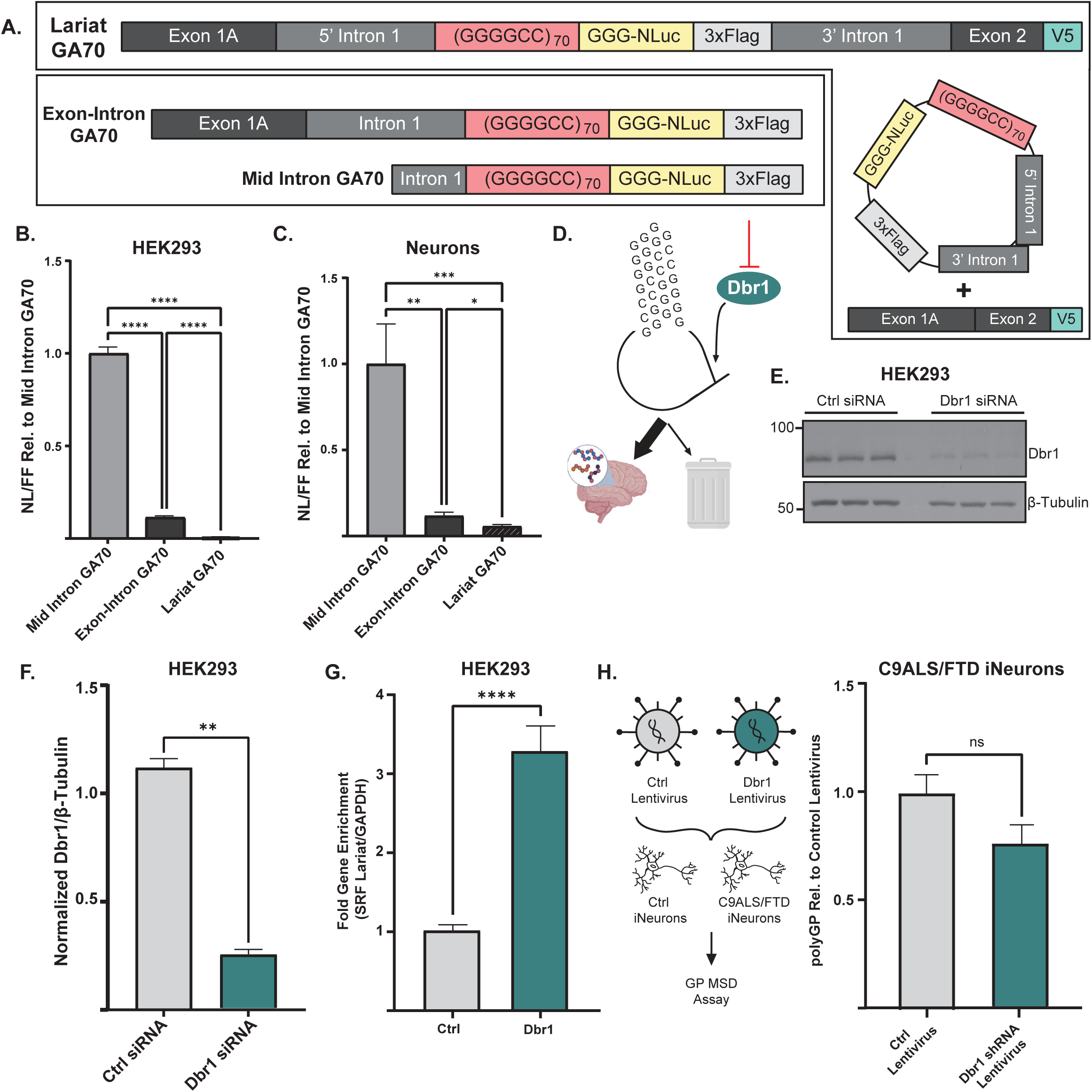
Novel linear intron-initiating repeat RNAs drive C9RAN translation. A) Schematic of mid intron GA70, exon-intron GA70, and C9 lariat (lariat GA70) nanoluciferase reporters. B) Expression of mid intron, exon-intron, and lariat GA70 reporters in HEK293 cells expressed as NL/FF relative to mid intron GA70 reporter, *n=9.* C) Expression of mid intron, exon-intron, and lariat GA70 reporters in rat hippocampal neurons expressed as NL/FF relative to mid intron GA70 reporter, *n=15.* D) Schematic of predicted Dbr1 impact a C9 lariat RNA. E) Western blot of Dbr1 protein levels in HEK293 cells transfected with non-targeting control or Dbr1 siRNA, 48 hours post-knockdown (KD). Tubulin was used as a loading control, *n=3*. F) Quantification of western blot (E) of Dbr1 proteins levels in HEK293 cells transfected with non-targeting control or Dbr1 siRNA, 48-hours post-KD. Expressed as relative intensity normalized to non-targeting control siRNA, *n=3*. G) Fold enrichment in gene expression of SRF lariat in HEK293 cells transfected with non-targeting control or Dbr1 siRNA, 48-hours post-KD. Normalized to GAPDH and expressed as relative to non-targeting control siRNA, *n=17.* H) Quantification of GP by MSD assay from C9ALS/FTD iNeurons treated with control or Dbr1 shRNA expressing lentivirus, expressed as arbitrary units relative to control shRNA lentivirus, *n=9*. Graphs represent mean ± SEM, * *p* < 0.05; *** *p* < 0.001; **** *p* < 0.0001. GA = glycine–alanine; GP = glycine-proline. (B-C) Brown-Forsythe one-way ANOVA with Welch’s unpaired t-test multiple comparison correction (F-H) Two-tailed student’s t-test with Welch’s correction.

To compare the translation efficiency of these reporters to the intron-initiated products, we transfected these linear and lariat GA70 reporters into HEK293 cells and rat hippocampal neurons. The mid-intron reporter corresponding to the most common intronic initiation site identified by 5’ RACE was by far the most efficiently expressed construct in both systems (**Figure 4B-C**). Consistent with prior studies in rabbit reticulocyte lysate (19), inclusion of exon1A impaired expression compared to intron-only GA70 reporters. In both HEK293 cells and rat hippocampal neurons, linear GA70 reporters were expressed between 10-to 100-fold more robustly than the lariat GA70 lariat reporter (**Figure 4B-C**). Despite this, NLuc signal of lariat GA70 was significantly greater than that of mock transfection (**Supplemental Figure 4B**), suggesting that the lariat reporter is engaging with translational machinery and generating products, consistent with previous results (22, 35).

Dbr1 is responsible for the cleavage of the 2’ to 5’ phosphodiester bond formed at the branch point of circularized lariats. Upon cleavage, the circular lariat becomes linearized, and due to its lack of protective 5’ cap or polyA tail, is rapidly degraded by exonucleases (45–47). We hypothesized that inhibition of Dbr1 would inhibit the cell’s ability to clear lariats, thus stabilizing these typically highly transient species, and that if a circularized lariat is the main driver of C9 RAN translation, expression of our lariat GA70 reporter would increase (**Figure 4D**). To assess this, we took advantage of an endogenous lariat RNA that is stable across multiple human cell types (34). The serum response factor (SRF) lariat is detectable in HEK293 cells at baseline (**Supplemental Figure 4C**).

Knockdown of Dbr1 by siRNA in HEK293 cells led to a roughly 4-fold enrichment the SRF lariat RNA compared to a non-targeting control siRNA (**Figure 4E-G**). As expected, knockdown of Dbr1 did not impact translation of linear GA70 reporters (**Supplemental Figure 4D**). However, Dbr1 reduction had no impact on lariat GA70 elicited NLuc expression, despite a 75% reduction in Dbr1 at the protein level (**Figure 4E-F, Supplemental Figure 4D**). While Dbr1 knockdown trended towards enhancement of lariat GA70 reporter-derived lariats as measured by RT-PCR across the branchpoint, this change was not significant due to high variance and low signal, suggesting that even reporter-based expanded repeat C9 lariats are difficult to stabilize in cells (**Supplemental Figure 4E**).

We next assessed whether we could measure circular C9 intron lariats endogenously in patient-derived iNeurons. RNA was extracted from iNeurons and treated with RNaseR, an RNase that selectively degrades linear RNA while leaving circular RNA intact. This methodology significantly enhanced our detection of the SRF lariat compared to mock treated patient-derived iNeurons and significantly reduced the levels of linear mature C9orf72 (**Supplemental Figure 4G**). However, we were unable to detect the presence of a circular C9 intron lariat despite allowing for 45 cycles of qRT-PCR. To confirm that our primers could bind their complementary sequence and generate a PCR product despite crossing a lariat branch point, we developed a reporter plasmid containing the unique branch point crossing sequence, which is only present when the intron is circularized in a lariat and arranged it linearly (linear C9 lariat). (**Supplemental Figure 4G**). PCR of the branch point crossing primers on the linear C9 lariat reporter was successful at inputs down to 0.005ng over 40 cycles (**Supplemental Figure 4G**). This linearly arranged sequence was readily detectable in transfected HEK293 cells (**Supplemental Figure 4H**). The lack of readily detectable C9 lariat is in-line with previous work aimed at measuring the circular C9 intron lariat, which required multiple rounds of 40-cycle PCR to identify the circular C9 intron boundary (35).

We next sought to investigate whether reducing Dbr1 levels in patient-derived iNeurons might impact endogenous DPR levels. We hypothesized that if an endogenous C9 intron lariat with a large repeat were to act as a critical endogenous RAN translation template, then Dbr1 knockdown should increase the levels of toxic DPRs (**Figure 4D**). We measured baseline polyGP levels in four C9 and four control patient-derived iNeuron lines, two of which were isogenic pairs (**Supplemental Figure 5A**). One line expressed significantly higher levels of polyGP compared to the others tested (10-fold or greater), which we used for our Dbr1 studies moving forward (**Supplemental Figure 5A**). Using a lentivirus expressing a Dbr1 shRNA, we transduced DIV3 C9 and control iNeurons and harvested cells at DIV10 to assess Dbr1 protein knockdown by western blot. Transduction with Dbr1 shRNA lentivirus reduced protein levels by roughly 50% in control and C9ALS/FTD iNeurons (**Supplemental Figure 5B-C**). We next measured polyGP levels in C9 iNeurons treated with a Dbr1 shRNA lentivirus or a control lentivirus and saw no significant change in polyGP levels in C9 iNeurons (**Figure 4H**). Taken together, these data suggest that a circular C9 intron lariat is unlikely to be an abundant or efficient template for RAN translation in patient-derived neurons.

## DISCUSSION

A central question in C9ALS/FTD pathogenesis is how an intronic repeat sequence is translated into toxic DPRs. Evidence suggests that RAN translation is important in disease pathogenesis and knowledge of its endogenous template in patient neurons could inform the development of therapeutics, such as antisense oligonucleotides (ASOs) and small molecules that target DPR production. Here we show that the repeats are found in linear, 5’ m^7^G-capped and sometimes polyadenylated mRNAs generated from transcriptional initiation within the intron itself in both a BAC transgenic mouse model of the disease and patient iNeurons. RAN translation of these cryptic mRNAs is highly efficient across all reading frames compared to lariat RNAs or retained exon-intron RNAs – both of which also appear to be less abundant. Based on these findings, we propose that these intron-initiating repeat containing linear RNAs are a major endogenous template for RAN translation in C9ALS/FTD, with important implications for therapy development.

We utilized a relatively new approach that couples nanopore sequencing with 5’ RACE as opposed to traditional molecular cloning followed by Sanger sequencing of individual clones (**Figure 1B**) (48). This methodology greatly increased both the throughput and sensitivity of the assay to detect low frequency transcripts, thus generating a more accurate view of the transcriptional landscape of the 5’ end of C9orf72. However, the absence of allele-specific SNPs within the region of interest precludes our ability to distinguish whether the transcripts we identified in our patient-derived iNeurons derive from the expanded disease allele or the non-expanded wildtype allele. While this cannot be resolved in our iNeuron model, the BAC transgenic C9 mouse model exclusively contains the expanded human allele, and insufficient sequence homology from the murine *C9orf72* gene prevents amplification by the *C9orf72*-specific 5’ RACE primers used in these experiments (**Supplemental Figure 1A**). Thus, intronic-initiated transcripts do occur in the presence of a large GGGGCC repeat (**Figure 1D-F**).

Identification of the same novel 5’ end transcripts by 5’ RACE of poly-adenylated mRNA suggests that, for at least a portion of these mRNAs, transcription proceeds through the repeat and to a polyA site (**Figure 2H-L**). While it is possible for pre-mRNA to be prematurely poly-adenylated, the stability of transcripts with this hallmark is markedly reduced. As such, our results are most consistent with the poly-adenylation of these novel transcripts occurring at non-premature sites (49). This is consistent with recent reports of cytoplasmic repeat-containing RNA mapping to C9orf72 exons and suggests that intron-initiated transcripts may be “exonized” and contain the expected downstream exons (38). Further, “exonized” transcripts are known to be actively translated, as siRNAs against exon 2 in C9ALS/FTD fibroblasts exhibits significantly decreased abundance of polyGA and polyGP DPRs (38).

Importantly, we observe 5’ intron-initiated mRNAs within polysomes from C9 patient-derived iNeurons, suggesting that these transcripts undergo translation in a disease relevant cell type and contribute to DPR production (**Figure 2A-B**). While our nanopore-coupled 5’ RACE technique greatly increased our ability to detect 5’ RACE products compared to traditional cloning and Sanger sequencing, we did not identify the shorter 32 nucleotide product seen in fibroblasts and iNeurons, likely due to a size-recovery limitation of PCR clean up. Over the past several years, long-read sequencing through repetitive regions has improved dramatically (50–53), and future 5’ RACE endeavors may be able to utilize primers that bind 3’ to the repeat to more reliably capture even very short products initiating just above the repeat.

In the classical scanning model of translational initiation, the 43S preinitiation complex made up of eIF2, GTP, Met-tRNAi^Met^, and 40S ribosome binds to capped mRNA through interaction with eIF4F, which includes the cap binding factor eIF4E (54). As such, steric hinderance induced by eIF4E is thought to preclude access of the 40S ribosome to the very 5’ end of capped mRNAs, creating a region of mRNA that is “blind” to the ribosome

(55). Despite this, using *in vitro* transcribed mRNA nanoluciferase reporters that mimic the novel 5’ end transcripts we identified through 5’ RACE, we observed robust translation from all three reading frames even when a very short leader was utilized that started just eight nucleotides above the CUG near-cognate codon previously shown to be important for polyGA reading frame translation (18, 19, 21, 25–27, 29, 44) (**Figure 3B-D, Supplementary Figure 2A-C**). Mutating this CUG codon to CCC markedly reduced production from this short intron reporter (**Figure 3G-H, Supplemental Figure 2D-E)**, and its translation was largely cap-dependent (**Figure 3I, Supplemental Figure 3A-E**). There are prior precedents for translation initiation at AUG codons within this typically “blind” region. Translation Initiator of Short 5’ UTR (TISU) is a short (twelve nucleotide median length) regulatory element located in 4.5% of protein-encoding genes. This element is typically close to the transcription start site plays important roles in the translation of mRNAs in the absence of ribosome scanning (56). Translation of *in vitro* transcribed GFP reporter TISU mRNAs remained, 7G cap-dependent and was not significantly altered when the 5’ UTR was shortened to only five nucleotides, consistent with our short intron GA70 reporter findings (**Figure 3I, Supplemental Figure 3A-E**). Similarly, histone H4 mRNAs also have short 5’ UTRs that utilize a hybrid cap-dependent scanning and IRES-driven initiation process (57). In this context, the histone ORF contains a highly structured helix region that directs positioning of the ribosome on the cognate start codon in the absence of ribosomal scanning (57). The GGGGCC repeat is known to adopt a strong secondary structure as a hairpin or g-quadruplex (58–61), suggesting it may act in a similar manner to support translation from a short leader sequence.

Our data suggest that the novel linear repeat-containing RNAs we observe in C9ALS/FTD patient neurons are much more efficient templates for RAN translation than either retained intronic repeat or spliced lariats that contain repeats. Previous data using a splice-capable C9 intron lariat reporter demonstrate that a spliced, circular intron can be exported to the cytoplasm and serves as the template for RAN translation (35). This work primarily utilized a reporter containing highly structured exogenous sequences (MS2 and PP7 binding sites) binding sites to track the mRNA, which could potentially influence the behavior of these reporters (62). Therefore, we sought to determine the contribution of a circular C9 intron lariat from a reporter lacking these sequences. Consistent with previous findings where the percentage of translating introns as measured by SunTag signal was markedly reduced compared to AUG-dependent translation (35), translation of our lariat reporter is inefficient in both HEK293 and rat hippocampal neurons (**Figure 4B-C**). This is likely due to inefficient, cap-independent translation mechanism by which a lariat would translate. Consistent with this theory, RAN translation of intron 1 is also inefficient when included as part of a bicistronic reporter system, where translation of a monocistronic reporter was ∼20-30 fold higher than with a bicistronic reporter (22).

We were unable to measure a repeat-containing intron lariat in C9 patient-derived iNeurons despite post-hoc enrichment for lariats by treating total RNA with RNaseR and assaying to 45 cycles by qRT-PCR (**Supplemental Figure 4F**). Importantly, this technique did significantly enrich for a known stable lariat (SRF2). Prior findings required deep sequencing following two rounds of PCR (35), suggesting that endogenous C9 intron lariats are maintained at a very low abundance within patient cells. To counter that limitation, we enriched for lariats by using a Dbr1 shRNA lentivirus in C9 patient-derived iNeurons.

However, this strategy had no impact on endogenous DPR production (**Figure 4H, Supplemental Figure 5B-D**). While Dbr1 knockdown did not impact polyGP production, future analysis of additional reading frames including polyGA and GR could reveal frame-specific effects.

In summary, our data demonstrates that intron-initiated repeat containing transcripts are more efficiently translated than either exon-intron or spliced lariat reporters and are more abundant in patient-derived iNeurons. These findings suggest that novel intron-initiated mRNAs represent the major endogenous RAN translation template in C9orf72 repeat associated FTD and ALS. This finding has significant therapeutic implications, as some, but not all, currently designed ASOs and other therapeutic approaches target this species. We propose that these cryptic intronic initiation sites could be viewed as unique therapeutic targets. Future studies will be needed to define what approach would be most effective.

## ACKNOWLEDGEMENTS

This work was supported by National Institutes of Health (F31127371 to SLM; F31NS100302 to KMG; R01NS097542, P01NS08497 and R35NS097273 to LP; R01NS113943 and 1R56NS128110-01 to SJB; R21HG011493 and R01GM144484 to APB; R01NS099280 to PKT; P30AG072931 to the University of Michigan Brain Bank and Alzheimer’s Disease Research Center), The Mayo Clinic Foundation to LP, Target ALS Foundation to LP, Robert Packard Center for ALS Research at Johns Hopkins University to LP, SJB and PKT, A. Alfred Taubman Medical Research Institute at the University of Michigan to APB and PKT, and the Veteran’s Administration (VA BLRD BX004842 to PKT).

We acknowledge support from the Bioinformatics Core of the University of Michigan Medical School’s Biomedical Research Core Facilities (RRID:SCR_019168), and from the Vector Core of the University of Michigan Medical School’s Biomedical Research Core Facilities for packaging lentivirus for this study.

## COMPETING INTERESTS

L.P. serves as a consultant for Expansion Therapeutics. P.K.T. serves as a consultant with Denali Therapeutics and has licensed technology through the University of Michigan to Denali. The remaining authors declare no competing financial interests.

## DATA, MATERIALS, AND SOFTWARE AVAILABILITY

Most data are included in the manuscript and supplemental appendices, including sequences of identified 5’ RACE products. Aligned BAMs are available upon request from the authors. Custom Python software used for nanopore based 5’ RACE transcript length determination using the Pysam module is available upon request.

## METHODS

### Cell Lines/Cell culture

**HEK293 cells** were purchased from American Type Culture Collection (ATCC). HEK293 cells were maintained in 10cm plates at 37°C, 5% CO_2_ in DMEM with high glucose (Gibco, 11965118) supplemented with 10% fetal bovine serum (50mL FBS added to 445mL DMEM; Bio-Techne, S11150) and 1% penicillin-streptomycin (5mL in in 445mL DMEM; Gibco, 15070063).

**Patient fibroblasts** were maintained in 10cm plates at 37°C, 5% CO_2_ in DMEM with high glucose supplemented with 10% fetal bovine serum, 1% MEM non-essential amino acids (Corning, MT25025CI) and 1% penicillin-streptomycin. When confluent, cells were pelleted at 200xg at room temperature for 5 minutes. Media was aspirated and cell pellets were flash frozen in liquid nitrogen. Pellets were stored at –80°C until use.

**iNeurons** (iNs) were differentiated according to (63). Briefly, induced pluripotent stem cells were washed in PBS and incubated in prewarmed accutase. TeSR-E8 media was added to the plate, cells were collected and pelleted, media aspirated, and the pellet resuspended in fresh TeSR-E8 media. Neuroprogenitors (NPs) were created by culturing cells for 2 days in TeSR-E8 plus 2mg/mL doxycycline. After this, cells were collected via accutase, counted and frozen in TeSR-E8 containing 10% DMSO. To begin each iN plating (Day 1), frozen NP cells were plated on PLO/laminin coated plates in TeSR-E8 containing Dox and Rock inhibitor. Day 2: media was changed to transition media containing N2 with 1x NEAA supplement, 10ng/mL BDNF, 10ng/nL NT3, 0.2ug/mL laminin, 2mg/mL doxycycline in half TeSR-E8 media, half DMEM F12. Day 3: media was changed to B27 media with 1x B27 supplement, 1x Glutamax supplement, 10ng/mL BDNF, 10ng/mL NT3, 0.2ug/mL laminin, and 1x Culture One in Neurobasal-A.

**Rat hippocampal neurons**: All animal use followed both National Institutes of Health (NIH) and University of Michigan Committee on Use and Care of Animals guidelines. Hippocampi were dissected from Sprague-Dawley rat pups of both sexes. Cells were papain dissociated as in (64). Hippocampal neurons were plated on poly(D-lysine)-coated 96-well culture plates and were maintained for at 37°C in 5% CO_2_ for 7 days in neuronal grown media (NGM; neurobasal medium A supplemented with 1xB27, 1xGlutamax).

### Tissue Harvest

Animal use followed NIH guidelines and in compliance with the University of Michigan Committee on Use and Care of Animals. Wild type (FVB/NJ, strain number 001800) and C9-500 BAC (FVB/NJ-Tg(C9orf72)500Lpwr/J, strain number 029099) mice were obtained from The Jackson Laboratory (Bar Harbor, ME). C9-500 BAC mice were developed and deposited by Laura Ranum (39).Animals were sacrificed at 6 weeks by isoflurane overdose followed by decapitation. Brains were removed and the cortex and cerebellum separated. Brain regions were snap frozen and stored at –80°C until used.

### *5’* Repeat-RLM-RACE

5’ RACE was performed using the GeneRacer kit (Invitrogen, L150201) according to kit protocol. Total RNA from HEK293 cells and iNeurons was harvested using the RNeasy Mini Kit (Qiagen, 74106) according to manufacturer protocol. RNA was extracted from 6-week-old mouse cerebellum using TRIzol (Thermo Fisher, 15596026) according to manufacturer protocol. For mouse experiments, 2-3ug of DNase-treated total RNA was used for the initial CIP reaction. For iNeuron experiments, 1ug of DNase-treated total RNA, 40ng of DNase-treated polyA captured RNA, or 0.2ug of monosome/polysome fraction DNase-treated RNA was used for the initial CIP reaction. For fibroblast experiments, ∼2.5-4.5ug of DNase-treated total RNA was used in the initial CIP reaction. 1pmol of C_4_G_2_x4 was spiked into the first step of cDNA synthesis, and cycling parameters were as follows (5 minutes at 25°C, 60 minutes at 55°C, 15 minutes at 70°C). Control reactions with beta-actin primers were performed as per kit protocol. All PCR reactions utilized Platinum PCR SuperMix High Fidelity (Thermo Fisher, 12532016) with 1uL cDNA and touchdown protocol recommended in the kit protocol. For P1 and P2 primers, annealing was performed at 68°C with 25s extension. For the P3 primer, annealing was performed at 66°C with 25s extension. PCR products were cleaned with DNA Clean and Concentrator Magbead Kit (Zymo Research) according to manufacturer instructions. Purified PCR products were used for nanopore long read sequencing.

### polyA Capture

DIV10 iNeurons were harvested with the RNeasy Mini Kit (Qiagen), treated with TURBO DNase (Thermo Fisher, AM2238), and clean and concentrated with the RNA Clean and Concentrate-25 kit (Zymo Research, R1018) as described above. polyA isolation was performed on 2ug of DNase-treated total RNA using the NEBNext Poly(A) mRNA Magnetic Isolation Module (New England Biolabs, E7490L) according to the manufacturer’s express protocol.

### ONT Library Preparation of Amplicons and Analysis

Sequencing libraries consisting of amplicons were prepared using the ONT Native Barcoding kit 24 V14 (SQK-NBD114.24, ONT) as described here with the following modifications. 10uL of each amplified product was phosphorylated using 0.5μL of T4 PNK (M0201S, NEB) and 1.5μL 10x T4 DNA ligase buffer (B0202S, NEB) and 3μL of molecular biology grade water in a total of 15μL.

The phosphorylated products were ligated in separate PCR tubes with different barcodes in a 20μL reaction with 0.5μL Native Barcode (NB01-24), 1μL T4 DNA ligase (M0202L, NEB), 1μL of T4 DNA ligase buffer and 2.5 μL of water for 20 minutes at RT. 2μL of 0.5M EDTA was added to each tube to stop the barcode ligation and all reactions were pooled into a single 1.5mL microcentrifuge tube. The barcoded reactions were incubated with 1.1X CleanNGS beads (CNGS005, Bulldog Bio) for 10 minutes at RT with rotation. The samples were then placed on a magnet and washed twice with 700μL of 80% ethanol. Following the washes, the pooled samples were eluted in 36μL of water, and 1μL was used to quantify the DNA concentration on the Qubit. The Native Adapter (NA) was ligated to the samples in a 50μL ligation reaction (5μL T4 DNA ligase, 12.5μL LNB, 5μL NA) and rotated for 20 minutes at RT. Next, 1.1X CleanNGS beads were added and incubated for an additional 10 minutes at RT with rotation. The library was placed on a magnet and the supernatant was removed followed by two 150μL washes with Small Fragment Buffer (SFB). After the final wash, the bead pellet was allowed to air dry for 30 seconds and the library was eluted in 16μL EB, and 1μL was used to quantify the DNA on the Qubit. The sequencing library was prepared with at least 500ng of barcoded-adapted sample, 15μL Sequencing Buffer (SB), 5μL Library Beads (LIB), and sequenced on a Flongle R10.4.1 flow cell following the ONT Flongle loading method. ONT sequencing was performed for up to 24 hours using ONT Flongle flowcells. The data were basecalled with Dorado version 7.6.7 using the dna_r10.4.1_e8.2_400bps_5khz_sup model and a minimum qscore of 10. Sequence alignment was performed using minimap2, aligning reads to the human reference assembly GRCh38. On-target reads for each barcode were filtered and counted by transcript length leveraging the Pysam module in custom Python software.

### Monosome/Polysome Collection in iNeurons

Polysome profiling was performed on DIV10 iNeurons as described in (65). Briefly, RNA from samples that eluted in the monosome and polysome fractions was isolated using TRIzol (Thermo Fisher, 15596026), according to manufacturer protocol. For monosome/polysome isolated iNeuron experiments, ∼0.2ug RNA was used as input for the initial 5’ RACE reaction.

### Plasmids

Vectors expressing AUG-nLuc and CrPV were described in (41).

**Linear Reporters:** Sequence between the G_4_C_2_ repeats and the 5’ end of nanoluciferase, excluding the Age1 site, was removed by Q5-Site Directed Mutagenesis kit (New England Biolabs, E0554S) in a 0-repeat version of the Intron-GA pcDNA3.1+ plasmid from (18). Q5 Site-Directed mutagenesis was used to remove intron 1 sequence to create the mid intron GA and short intron-GA reporters, respectively. 70 G_4_C_2_ repeats were inserted into pcDNA3.1+ using NheI and AgeI. Site directed mutagenesis to adjust the reading frame of nanoluciferase from the GA to GP or GR frame was performed using Q5 Site-Directed Mutagenesis. Q5 Site-Directed Mutagenesis was used to mutate the CUG start codon to CCC in the short intron constructs. All ligations were performed with Rapid DNA Ligation Kit (Roche, 11635379001) with incubation at room temperature for 30 minutes. Repeat length was verified by restriction digest and gel electrophoresis.

**Lariat GA70:** A gene block containing exon1A through but not including the repeat in intron 1, GGG-nanoluciferase-3xFlag in the GA frame, the terminal 200 nucleotides of the 3’ end of intron 1, and the 5’ end of exon 2 upstream of the ATG start codon with a terminal V5 tag was ordered from Azenta. This gene block was inserted into pcDNA3.1+ using PspOMI and Acc65. 70 G_4_C_2_ repeats were excised from the Intron1-GA70 construct in K.M. Green, 2017 and inserted into pcDNA3.1+ using Asc1 and Eag1. All ligations were performed with T4 DNA ligase (Roche, 10481220001) with incubation at 4°C for 16 hours. Repeat length was verified by restriction digest and gel electrophoresis.

### RNA Synthesis

RNAs were in vitro transcribed from linearized plasmids as described in (18). For capping, two types of 5’ caps were added: 3’-O-Me-m7GpppG anti-reverse analog (ARCA) cap was used to functionally cap RNA (New England Biolabs, S1411S) or 3’-O-Me-m7AppG (A-cap) to protect RNA but preclude cap-dependent translation (New England Biolabs, S1406S). 500ng of *in vitro* transcribed RNA was assessed for size and quality on an agarose gel according to the Denaturation and Electrophoresis of RNA with Glyoxal protocol in (66).

### Rabbit Reticulocyte Lysate In Vitro Translation

Performed as in (18). Briefly, 3nM *of in vitro* transcribed RNA were translated with Flexi Rabbit Reticulocyte Lysate System that is supplemented with calf liver tRNA (Promega, L4540). Reactions for luminescence assays were programed with 3nM RNA and contained 30% RRL, 10mM amino acid mix minus methionine, 10mM amino acid mix minus leucine, 100mM KCl. 0.5mM MgOAc, and 0.8U uL-1 Murine RNase Inhibitor (New England Biolabs, M0314L). Reactions were incubated at 30°C for 30 minutes before termination by incubation at 4°C. 24 hours later, nanoluciferase assay was performed as described below.

### RNA Transfections

**HEK293 cells** were plated at 2×10^4^ cells per well in a 96 well plate. 24 hours later when cells were 60-70% confluent, 90ng of *in vitro* transcribed mRNA and 200ng of pGL4.13 firefly luciferase plasmid was transfected in triplicate using TransIT-mRNA Transfection Kit (Mirus, 2225) according to manufacturer protocol. 24 hours later, nanoluciferase assay was performed as described below.

**Rat hippocampal neurons** were plated at 3.5×10^4^ cells per well in a 96-well plate. 7 days later, cells were transfected with 50ng of nanoluciferase mRNA and 50ng of pGL4.13 firefly luciferase plasmid in 0.3uL Lipofectamine MessengerMAX reagent according to manufacturer protocol (Thermo Fisher, LMRNA008). 24 hours later, nanoluciferase assay was performed as described below.

### siRNA Transfection

0.5pmol of Dbr1 siRNA (Fisher, 4427037) or a non-targeting control siRNA (Thermo Fisher, 4390843) was combined with 0.16uL Lipofectamine^TM^ RNAiMAX Transfection Reagent (Thermo Fisher, 13778150) in Opti-MEM^TM^ Reduced Serum Media (Thermo Fisher, 31985070) and incubated at room temperature for 10 minutes. HEK293 cells were plated at 1.2×10^4^ cells per well in a 96 well plate following addition of the siRNA mixture to the 96 well plate. 48 hours later when cells were 60-70% confluent, 50ng of nanoluciferase reporter plasmid and 50ng of pGL4.13 firefly plasmid was mixed 3:1 with FuGENE HD Transfection Reagent (Promega, E2311) to DNA in Opti-MEM media and incubated at room temperature for 10 minutes prior to adding to cells. 24 hours later, nanoluciferase assay was performed as described above.

### Plasmid Transfections

**HEK293 cells** were plated at 1.6×10^4^ cells per well in a 96-well plate. 24 hours later when cells were 60-70% confluent, 50ng of nanoluciferase plasmid and 50ng of pGL4.13 firefly luciferase plasmid were mixed 3:1 with FuGENE HD transfection reagent to DNA in Opti-MEM media and incubated at room temperature for 10 minutes prior to adding to cells. 24 hours later, nanoluciferase assay was performed as described below.

**Rat hippocampal neurons** were plated at 3.5×10^4^ cells per well in a 96-well plate. 7 days later, cells were transfected with 50ng of nanoluciferase plasmid and 50ng of pGL4.13 firefly luciferase plasmid in 0.3uL Lipofectamine MessengerMAX reagent according to manufacturer protocol (Thermo Fisher, LMRNA008). 24 hours later, nanoluciferase assay was performed as described below.

### Luminescent Assays

**Rabbit Reticulocyte Lysate:** Reactions were diluted 1:7 in Glo Lysis Buffer (Promega, E2661) and incubated 1:1 with freshly prepared NanoGlo Substrate diluted 1:50 in NanoGlo Buffer (Promega, N1150) in the dark in opaque 96-well plates for 5 minutes at room temperature. Luminescence was then measured on a GloMax 96 Microplate Luminometer.

**HEK293 and Rat Hippocampal Neurons:** Cells were lysed in 70uL Glo Lysis Buffer 24 hours after transfection (Promega, E26661) and incubated 1:1 with freshly prepared NanoGlo Substrate diluted 1:50 in NanoGlo Buffer (Promega, N1150) on a plate rocker in the dark for 5 minutes. Cell lysate was incubated 1:1 in diluted NanoGlo Substrate on a plate rocker in the dark for 5 minutes to assess nanoluciferase or incubated 1: 1 with One-Glo on a plate rocker in the dark for 5 minutes to assess firefly luciferase using Luciferase Assay System (Promega, E6130). Luminescence was measured on a GloMax 96 Microplate Luminometer.

### Western Blot

**HEK293 Cells**: 10pmol of Dbr1 siRNA (Thermo Fisher, 4427037) or non-targeting control siRNA (Thermo Fisher, 4390843) was combined with 3.2uL Lipofectamine^TM^ RNAiMAX Transfection Reagent (Thermo Fisher, 13778150) in Opti-MEM^TM^ Reduced Serum Media (Thermo Fisher, 31985070) and incubated at room temperature for 10 minutes. HEK293 cells were plated at 2.8×10^5^ cells per well in a 12-well plate following addition of the siRNA mixture to the plate. 48 hours later when cells were ∼70% confluent, media was removed, and cells were washed 1x in ice cold phosphate buffered saline (PBS). PBS was removed and cells were lysed with 300uL ice cold RIPA buffer with cOmplete Mini protease inhibitor tablet (Roche, 11836153001) for 30 minutes at 4°C. Lysates were homogenized using a Series 60 Sonic Dismembrator (Thermo Fisher, discontinued) for 25 seconds. Lysates were diluted 1:200 in Bradford assay reagent and read at 595nm to quantify protein content in each sample. Lysates were denatured in 6x sample loading dye mixed 25:3 with 2-mercaptoethanol and boiled at 95°C for 5 minutes. Samples were stored at –20°C prior to use. 20ug sample was run on an 8% SDS-PAGE gel and transferred to PVDF membranes overnight at 30 V at 4°C. Membranes were blocked in 5% non-fat dry milk, and all antibodies were diluted in 5% non-fat dry milk. Washes were performed with 1x TBST 3 times, 8 minutes each. Dbr1 antibody was applied at 1:1,000 and incubated for 1.5 hours at room temperature. HRP anti-rabbit secondary antibody was applied at 1:10,000 and incubated for 1 hour at room temperature. Beta-tubulin was applied 1:1,000 and incubated for 1 hour at room temperature. HRP anti-mouse secondary antibody was applied at 1:10,000 and incubated at room temperature for 1 hour. Bands were visualized on film and band intensities were measured using ImageJ and quantified by band intensity with background subtracted, normalizing to beta-tubulin.

**iNeurons:** Media from DIV10 iNeurons was removed and cells were lysed with 300uL ice cold RIPA buffer with cOmplete Mini protease inhibitor tablet (Roche, 11836153001) for 30 minutes at 4°C. Lysates were homogenized using a Series 60 Sonic Dismembrator (Thermo Fisher, discontinued) for 25 seconds. Lysates were diluted 1:200 in Bradford assay reagent and read at 595nm to quantify protein content in each sample. Lysates were denatured in 6x sample loading dye mixed 25:3 with 2-mercaptoethanol and boiled at 95°C for 5 minutes. Samples were stored at –20°C prior to use. 40ug sample was run on an 8% SDS-PAGE gel and transferred to PVDF membranes overnight at 30 V at 4°C. Membranes were blocked in 5% non-fat dry milk, and all antibodies were diluted in 5% non-fat dry milk.

Washes were performed with 1x TBST 3 times, 8 minutes each. Dbr1 antibody was applied at 1:1,000 and incubated for 1 hour at room temperature. HRP anti-rabbit secondary antibody was applied at 1:10,000 and incubated for 1 hour at room temperature. Beta-tubulin was applied 1:1,000 and incubated overnight at 4°C. HRP anti-mouse secondary antibody was applied at 1:10,000 and incubated for 1 hour at room temperature. Bands were visualized on film and band intensities were measured using ImageJ and quantified by band intensity with background subtracted, normalizing to beta-tubulin.

### RNaseR

HEK293 and DIV10 iNeurons were harvested using RNeasy Mini Kit (Qiagen, 74106) according to manufacturer protocol. Up to 10ug of RNA was treated with TURBO DNase (Thermo Fisher, AM2238) according to manufacturer protocol and clean and concentrated with RNA Clean and Concentrator-5 (Zymo Research, R1016). DNase treated RNA was treated with 15U RNaseR (Molecular Cloning Laboratories, RNASR-200) by incubating at 37°C for 15 minutes. An equivalent amount of RNA was also mock treated with water at 37°C for 15 minutes. RNaseR treated RNA was clean and concentrated with RNA Clean and Concentrator-5 (Zymo Research, R1016).

### cDNA Synthesis

RNA was extracted from HEK293 cells and iNeurons as described above. Up to 10ug of RNA was treated with TURBO DNase (Thermo Fisher, AM2238) according to manufacturer protocol and clean and concentrated with RNA Clean and Concentrator-5 (Zymo Research, R1016). 0.5-1.5ug of cleaned RNA was reverse transcribed to cDNA using Superscript III according to manufacturer protocol (Thermo Fisher, 18080051).

### Quantitative Real-Time Reverse Transcription PCR (qRT-PCR)

RNA from HEK293 and iNeurons was isolated as described above. cDNA from each sample was generated from 250ng-1ug of RNA as described above. cDNA was diluted to 100ng for samples and a 3-point standard curve was generated at 2x, 0.2x, and 0.02x concentration. TaqMan primer-probe pairs are listed in Table X. TaqMan Fast Advanced Master Mix for qPCR (Thermo Fisher, 4444557) and fast cycling parameters (20s at 95°C, 1s at 95°C, 20s at 60°C) were used. cDNA abundance was measured after 40 cycles unless otherwise indicated using QuantStudio3 (Applied Biosystems) and calculated using ΔΔCT method.

Experiments were performed in triplicate, with 3 wells per experiment.

### Lentivirus

A lentiviral shRNA plasmid against Dbr1 was purchased from Horizon Discovery. GFP control vector (pLLEV-GFP) was purchased from the University of Michigan Vector Core. Lentiviruses were packed at the University of Michigan Vector Core with HIV lentiviruses and then concentrated to 10x concentration in 10mL of DMEM. Knockdown of Dbr1 was confirmed by western blot. To transduce iNeurons, media was removed from control and C9 iNeurons on DIV3, and a full media change with diluted control GFP lentivirus or Dbr1 shRNA lentivirus in B27 was applied. A half media change with B27 media was performed on DIV6, and iNeurons were harvested on DIV10 according to the protocol detailed below for GP MSD.

### GP MSD Assay

From a 6-well plate, media from DIV10 iNeurons was removed manually, and cells were harvested by scraping with 200uL cold co-IP buffer (50mM Tris-HCl, 300mM NaCl, 5mM EDTA, 0.1% triton-X 100, 2% SDS, protease inhibitor, phosphoSTOP) on ice. Lysates were passed through a 21G syringe 12 times, spun at 16,000xg for 20 minutes at 15°C, and supernatant was collected. polyGP protein levels were measured using the Meso Scale Discovery (MSD) electrochemiluminescence detection technology as previously described (67). Detection was performed as described in (65)

---

**Supplemental Figure 1:**
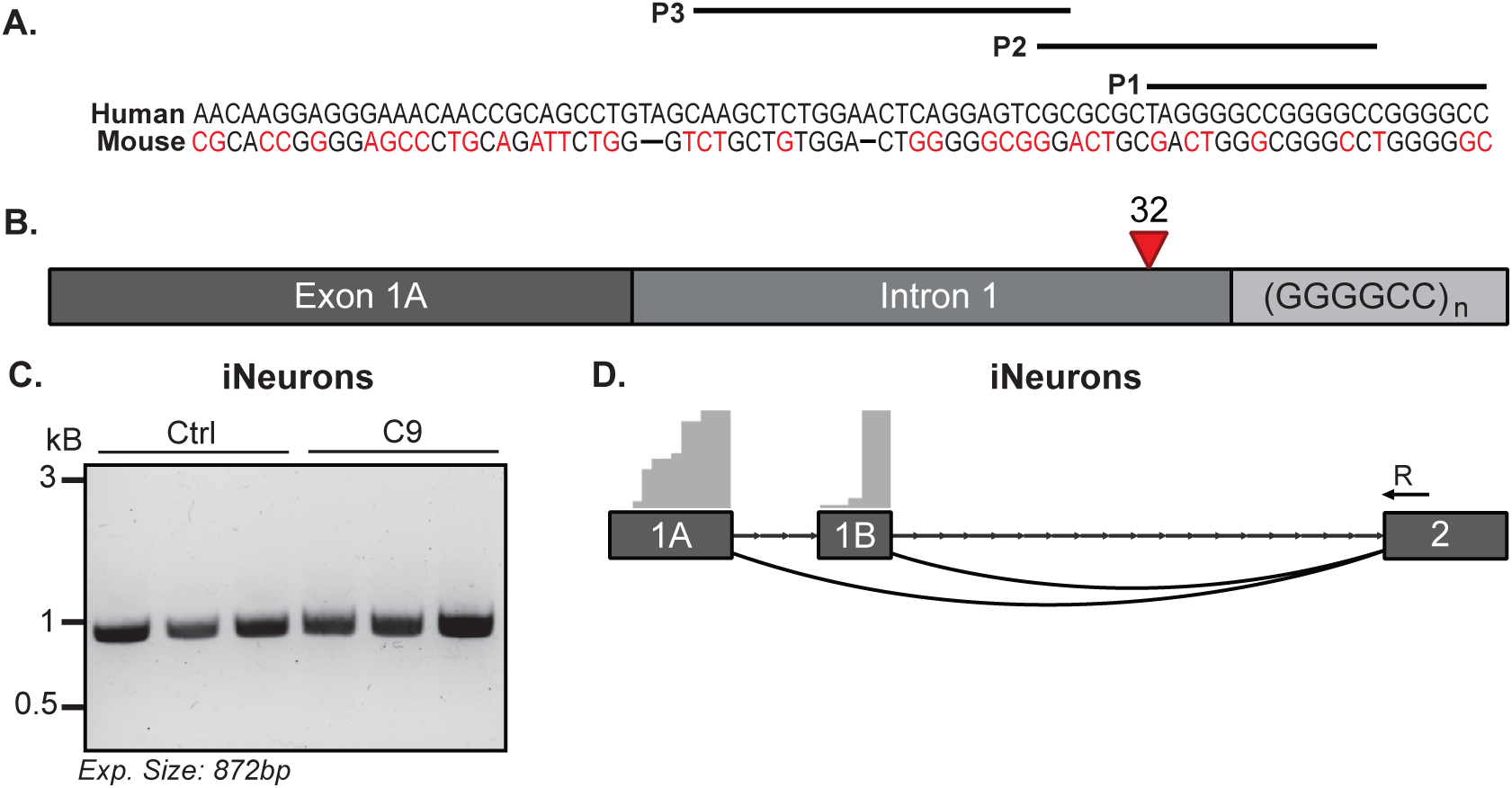
Repeat RLM-RACE reveals novel repeat-containing RNA templates in C9-500 BAC mice and patient derived cells (Relevant to Figures 1 and 2). A) Sequence diferences of C9orf72 between human and mouse and sites of Primers P1-P3 used in RACE studies showing lack of alignment to mouse sequence. B) Schematic of the most commonly identified 5’ RACE products using primers 2 and 3 in two C9ALS/FTD and 1 control fibroblast lines. C) Agarose gel of human Beta-actin control 5’ RACE PCR in C9ALS/FTD and control iNeurons, *n=3.* D) Aligned IGV plot using an exon 2 reverse primer in total RNA from C9ALS/FTD and control iNeurons, *n=2/group*.

**Supplemental Figure 2:**
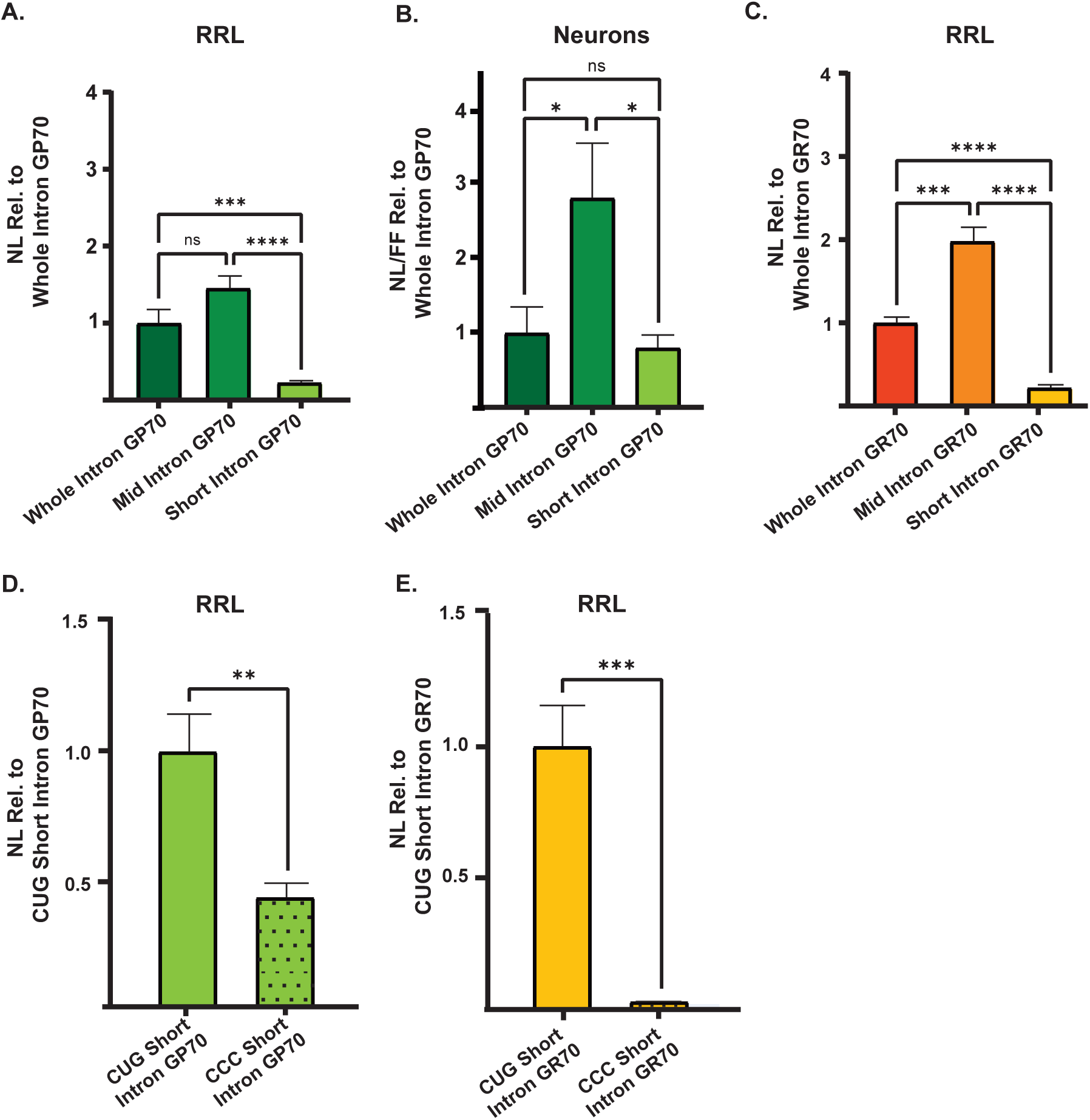
Cryptic intron-initiating mRNAs are robustly translated across model systems (relevant to figure 3). A) Expression of short, mid, and whole intron GP70 reporters in RRL, expressed NL relative to whole intron GA70, *n=9*. B) Expression of short, mid-length, and whole intron GP70 in rat hippocampal neurons, expressed as NL/FF relative to whole intron GP70, *n=15.* C) Expression of mid-intron initiating and whole intron GR70 reporters in RRL, expressed as NL relative to whole intron GR70, *n=9*. D) Expression of short-intronic initiating GP70 reporters in RRL, expressed as NL relative to CUG short intron GP70. polyGA frame CUG was mutated to CCC, *n=9*. E) Expression of short-intronic initiating GR70 reporters in RRL, expressed as NL relative to CUG short intron GR70. polyGA frame CUG was mutated to CCC, *n=9.* Graphs represent mean ± SEM, * *p* < 0.05; *** *p* < 0.001; **** *p* < 0.0001. GP = glycine–proline; GR = glycine–arginine; CrPV = cricket paralysis virus. (A-E) Two-tailed student’s T-test with Welch’s correction.

**Supplemental Fig. 3:**
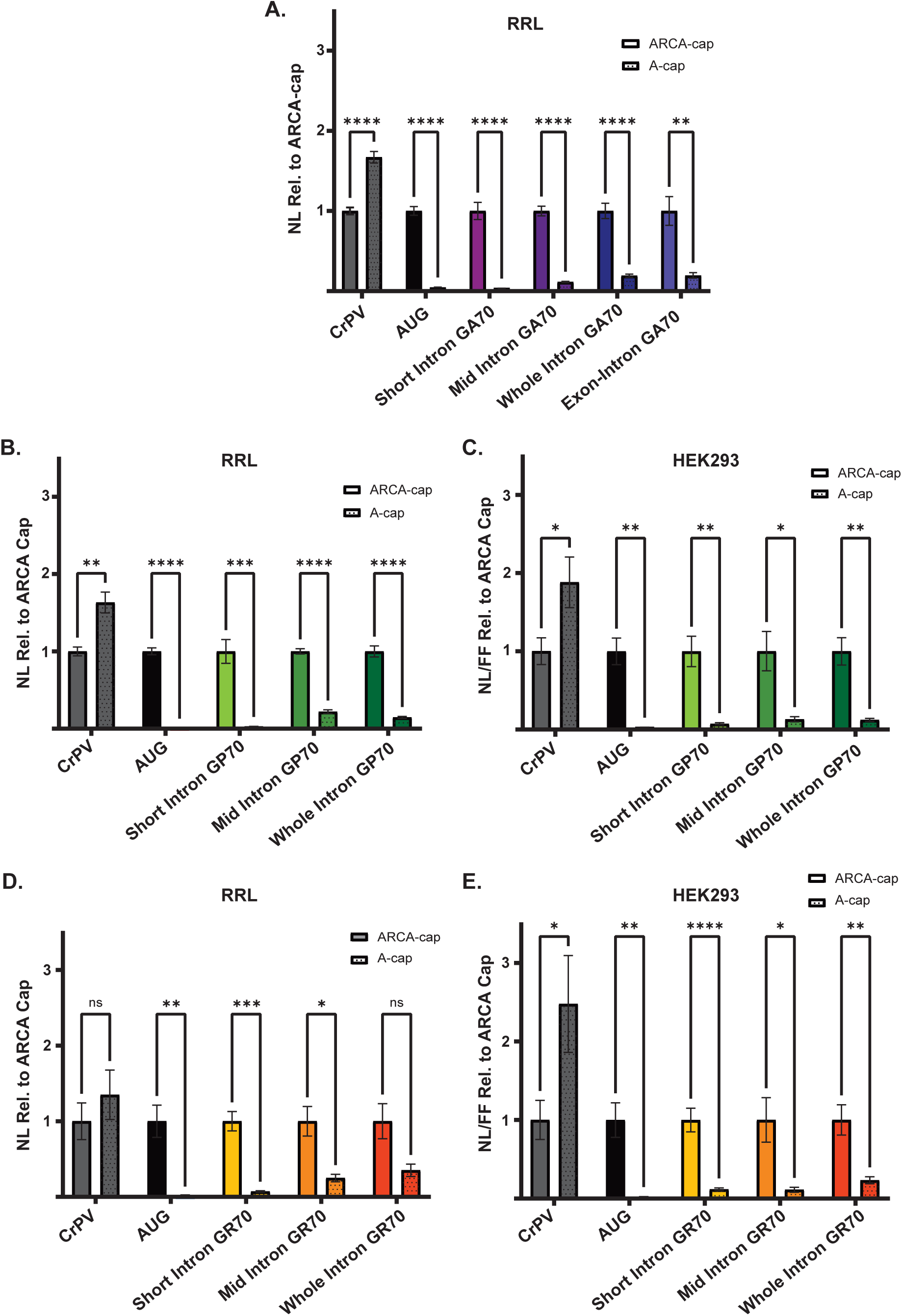
Novel intronic initiating reporter RNAs are translated in a cap dependent manner and more robustly than intron inclusion RNAs (Relevant to figure 3). A) Expression of ARCA m^7^G-capped and A-capped control short intron, mid intron, whole intron, and exon-intron GA70 reporters in RRL, expressed as the ratio of NL/FF signal in A-capped reporters to ARCA m^7^G-capped reporters, *n=9*. B) Expression of ARCA m^7^G-capped and A-capped control short, mid, and whole intron GP70 reporters in RRL, expressed as the ratio of NL/FF signal in A-capped reporters to ARCA m^7^G-capped reporters, *n=9.* C) Expression of ARCA m^7^G-capped and A-capped control short, mid, and whole intron GP70 reporters in HEK293 cells, expressed as the ratio of NL/FF signal in A-capped reporters to ARCA m^7^G-capped reporters, *n=9*. D) Expression of ARCA m^7^G-capped and A-capped control short, mid, and whole intron GR70 reporters in RRL, expressed as the ratio of NL/FF signal in A-capped reporters to ARCA m^7^G-capped reporters, *n=9*. E) Expression of ARCA m^7^G-capped and A-capped short, mid, and whole intron GR70 reporters in HEK293 cells, expressed as the ratio of NL/FF signal in A-capped reporters to ARCA m^7^G-capped reporters, *n=9*. Graphs represent mean ± SEM, * *p* < 0.05; *** *p* < 0.001; **** *p* < 0.0001. GP = glycine–proline; GR = glycine–arginine, CrPV cricket paralysis virus. (A-E) Multiple Holm-Šídák two-tailed student’s T-test with Welch’s correction.

**Supplemental Fig. 4:**
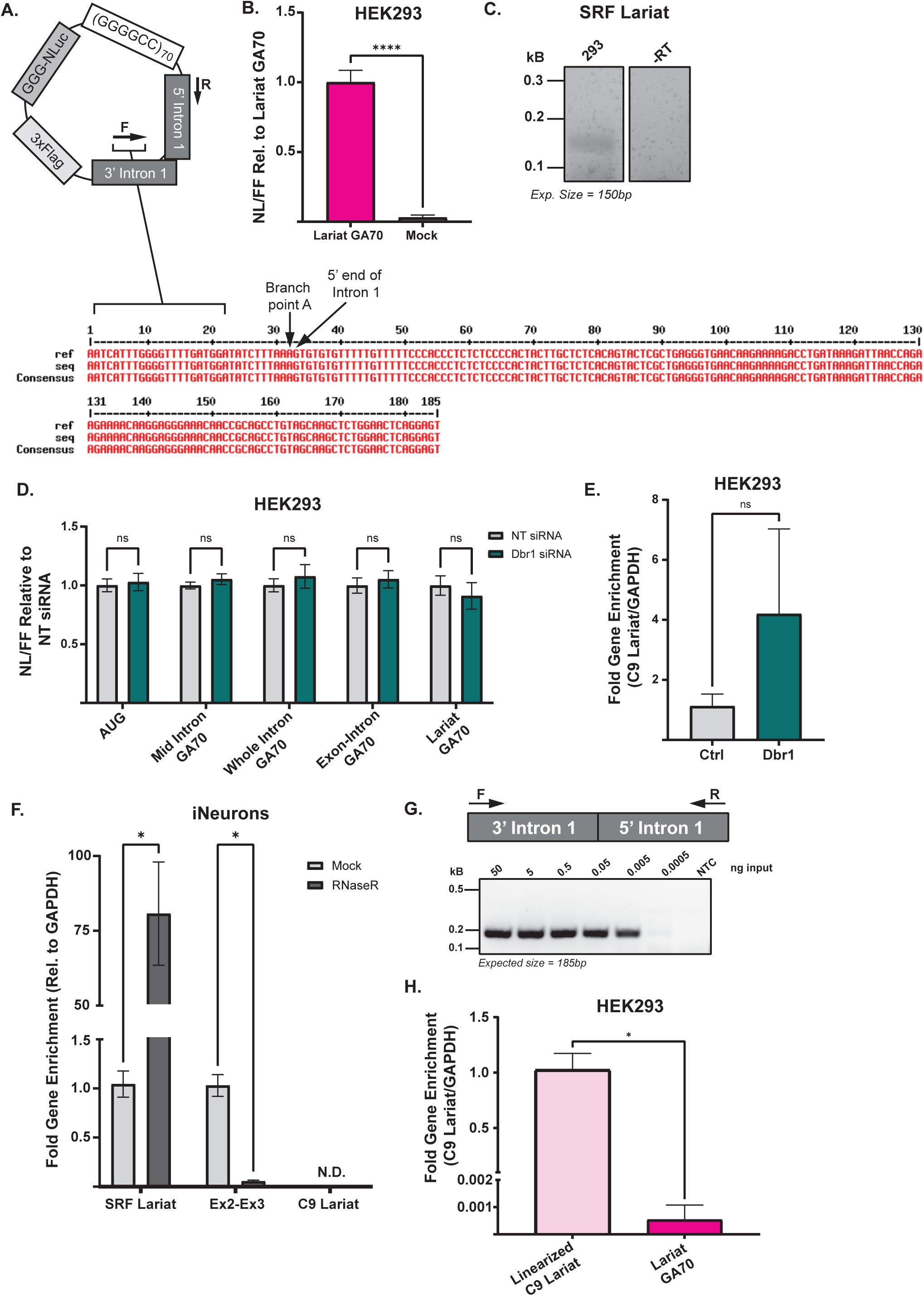
C9 repeat lariat detection and enrichment. A) Sequencing results of transfected lariat GA70 reporter in HEK293 cells. B) Expression of lariat GA70 reporter compared to mock transfection in HEK293 cells, expressed as NL/FF relative to lariat GA70 reporter, *n=9*. C) Agarose gel of SRF lariat PCR across the branch point in 293 cells. D) Expression of mid intron, whole intron, exon-intron, and lariat GA70 reporters in HEK293 cells, expressed as NL/FF ratio relative to non-targeting control siRNA, *n=9*. E) Fold gene enrichment of C9 lariat in HEK293 cells transfected with non-targeting control or Dbr1 siRNA, 48-hours post-KD. Normalized to GAPDH and expressed as relative to non-targeting control siRNA, *n=3*. F) Fold gene enrichment of SRF lariat, Exon 2-Exon 3 C9orf72, and C9 lariat in mock treated or RNaseR treated patient-derived iNeurons. Normalized to GAPDH and expressed as relative to mock treated iNeurons, *n=6*. G) Schematic of primer binding on C9 lariat reporter and agarose gel of linearized C9 lariat reporter PCR product. H) Fold gene enrichment of transfected lariat reporters, measured with branch point crossing primers. Normalized to GAPDH and expressed as relative to linearized C9 lariat, *n=3*. Graphs represent mean ± SEM, * *p* < 0.05; *** *p* < 0.001; **** *p* < 0.0001. GA = glycine-alanine. (B, E, H) Two-tailed student’s T-test with Welch’s correction, (D, F) Multiple Holm-Šídák two-tailed student’s T-test with Welch’s correction.

**Supplemental Fig. 5:**
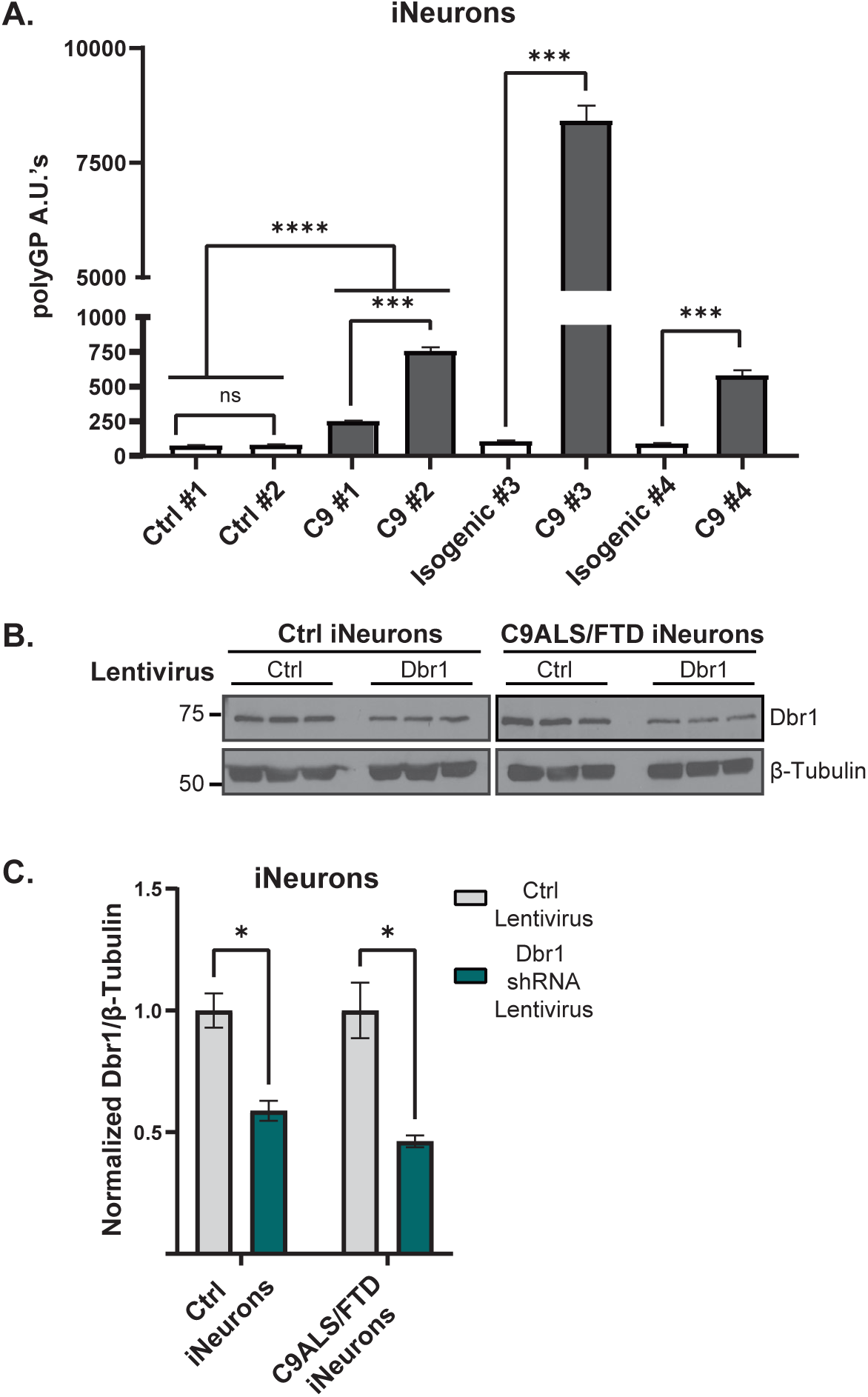
Dbr1 knockdown in patient derived iNeurons does not reduce levels of endogenous DPRs. A) Quantification of poly GP by MSD assay from control and C9ALS/FTD iNeurons, expressed as arbitrary units, *n=3.* B) Western blot of Dbr1 protein levels in C9ALS/FTD and control line used for GP MSD experiment following Dbr1 knockdown (KD). Tubulin was used as a loading control, *n=3.* C) Quantification of western blot (B) of Dbr1 proteins levels in control and C9ALS/FTD iNeurons transduced with control or Dbr1 shRNA expressing lentivirus, 7-days post-KD. Expressed as relative intensity normalized to non-targeting control shRNA lentivirus, *n=3.* Graphs represent mean ± SEM, \**p* < 0.05; *** *p* < 0.001; **** *p* < 0.0001. GP – glycine-proline. (A, C) Multiple Holm-Šídák two-tailed student’s T-test with Welch’s correction.

## Notes

### Competing Interest Statement

The authors have declared no competing interest.

